# A Cross-Species Atlas of the Dorsal Vagal Complex Reveals Neural Mediators of Cagrilintide’s Effects on Energy Balance

**DOI:** 10.1101/2025.01.13.632726

**Authors:** Mette Q. Ludwig, Bernd Coester, Desiree Gordian, Shad Hassan, Abigail J. Tomlinson, Mouhamadoul Habib Toure, Oliver P. Christensen, Anja Moltke-Prehn, Jenny M. Brown, Dylan M. Rausch, Anika Gowda, Iris Wu, Stace Kernodle, Victoria Dong, Mike Ayensu-Mensah, Paul V. Sabatini, Jae Hoon Shin, Melissa Kirigiti, Kristoffer L. Egerod, Christelle Le Foll, Sofia Lundh, Marina Kjærgaard Gerstenberg, Thomas A. Lutz, Paul Kievit, Anna Secher, Kirsten Raun, Martin G. Myers, Tune H. Pers

**Affiliations:** Novo Nordisk Foundation Center for Basic Metabolic Research, University of Copenhagen, Copenhagen, Denmark; Digital Science & Innovation, Novo Nordisk A/S, Måløv, Denmark; Departments of Internal Medicine, University of Michigan and Molecular and Integrative Physiology, Ann Arbor, Michigan, USA; Global Drug Discovery, Novo Nordisk A/S, Måløv, Denmark; Oregon National Primate Research Center, Oregon Health & Science University, Beaverton, Oregon, USA; University of Zurich, Zurich, Switzerland

## Abstract

Amylin analogs, including potential anti-obesity therapies like cagrilintide, act on neurons in the brainstem dorsal vagal complex (DVC) that express calcitonin receptors (CALCR). These receptors, often combined with receptor activity-modifying proteins (RAMPs), mediate the suppression of food intake and body weight. To understand the molecular and neural mechanisms of cagrilintide action, we used single-nucleus RNA sequencing to define 89 cell populations across the rat, mouse, and non-human primate caudal brainstem. We then integrated spatial profiling to reveal neuron distribution in the rat DVC. Furthermore, we compared the acute and long-term transcriptional responses to cagrilintide across DVC neurons of rats, which exhibit strong cagrilintide responsiveness, and mice, which respond poorly to cagrilintide over the long term. We found that cagrilintide promoted long-term transcriptional changes, including increased prolactin releasing hormone (*Prlh*) expression, in the nucleus of the solitary tract (NTS) *Calcr/Prlh* cells in rats, but not in mice, suggesting the importance of NTS *Calcr/Prlh* cells for sustained weight loss. Indeed, activating rat area postrema *Calcr* cells briefly reduced food intake but failed to decrease food intake or body weight over the long term. Overall, these results not only provide a cross-species and spatial atlas of DVC cell populations but also define the molecular and neural mediators of acute and long-term cagrilintide action.

## Introduction

Amylin analogs reduce food intake and represent a promising anti-obesity therapy ^1^. Treatment with the long-acting amylin analog cagrilintide ^2^ induces 10.8% weight loss in humans over 26 weeks ^3^ and co-administering cagrilintide with the glucagon-like peptide-1 (GLP-1) receptor agonist (GLP1RA) semaglutide produces a 17.1% weight loss over 20 weeks (versus 9.8% for semaglutide alone) ^4^.

Endogenous amylin is co-secreted with insulin from pancreatic beta cells. It mediates its appetite-suppressing effects via the amylin receptor (Amy3R-a complex of CALCR and a receptor activity modifying protein (RAMP)), especially in neuron populations of the brainstem dorsal vagal complex (DVC) ^5–8^. The DVC consists of the area postrema (AP; a circumventricular organ that lies outside of the blood-brain barrier), the nucleus of the solitary tract (NTS; the major termination site of vagal afferents), and the dorsal motor nucleus of the vagus (DMV; responsible for parasympathetic outflow to peripheral organs including those of the gut) ^9^.

DVC *Calcr* neurons contribute to the control of food intake and body weight. Artificial activation of rodent AP and NTS *Calcr* neurons acutely reduces food intake and body weight ^10,11^ while lesioning the AP reduces the acute suppression of food intake induced by amylin ^5^. Moreover, knocking down DVC *Calcr* attenuates the amylin-mediated activation of DVC neurons and increases meal duration ^7^. This is consistent with our previous finding showing *Calcr*-expressing DVC neurons in the mouse express transcriptional programs preferentially encoded by genes located in obesity-associated genome-wide association study (GWAS) loci ^11^. These DVC *Calcr* neurons represent likely candidates for mediating the weight-reducing effects of amylin and its analogs.

Interestingly, whereas amylin analogs like cagrilintide mediate sustained weight loss in rats, they suppress feeding less robustly and only transiently in mice ^6,12–14^. While we and others have begun to molecularly identify cell populations in the mouse DVC ^11,15,16^, the molecular anatomy of this region remains incompletely defined. The conservation of DVC cell types across species is unknown, and the nature of the DVC mediators and mechanisms of action for amylin analogs remain unclear.

To improve the ability to understand brain function and mechanism of action of anti-obesity therapies, we generated a comprehensive mouse, rat, and non-human primate cross-species single-nucleus RNA sequencing (snRNA-seq) DVC atlas covering >581,630 cells across 89 cell populations. Subsequently we used spatial profiling of >98,998 cells to resolve the anatomical location of 68 neuronal cell populations. Using in vivo pharmacology, we found that while cagrilintide rapidly modulated gene expression in AP *Calcr* neurons in mice and rats, the long-term activation of prolactin-releasing hormone (*Prlh*) in NTS *Calcr* neurons was specific to the rat, suggesting a key role for NTS *Prlh* in the long-term response to cagrilintide. Indeed, while activating rat AP *Calcr* neurons suppressed feeding only transiently, NTS *Prlh* signaling was required for the long-term suppression of food intake and body weight by cagrilintide in rats.

## Results

### Cross-species transcriptional atlas of the DVC

To identify species-specific transcriptional states underlying cagrilintide signaling in the DVC, we treated 30 diet-induced obese (DIO) mice and 42 DIO rats with either a single dose or seven daily subcutaneous (s.c.) treatments of cagrilintide or vehicle. To enable examination of transcriptional effects independent of weight loss, six of the vehicle-dosed mice and 11 of the vehicle-dosed rats were weight-matched to the cagrilintide-dosed animals. From these animals, we isolated the DVC-containing caudal brainstem and subjected this tissue to snRNA-seq (**Fig. 1a, Supplementary Table 1**). Moreover, we included six lean mice and rats in the single-cell atlas, as well as 12 mice treated with either acute or subchronic injections of the Amy3R-selective amylin analog AM1213 ^17^. The lean control animals were included to identify transcriptional changes induced by DIO. We also applied snRNA-seq to the DVC of five normal weight non-human primates (rhesus macaque) to assess conservation of cell populations and receptor profiles between rodents and a species more similar to humans. We split the atlases from each species into glial cells and neurons, mapped genes in rats and rhesus macaques to mouse orthologous genes and integrated the glial cells and neurons across species. To further increase the coverage of cell populations from the AP and to increase confidence in the clustering, we also included cells used to generate our previous AP-centric mouse atlas ^11^. In total these studies yielded an atlas of 319,447 glial cells and 262,226 neurons (**Supplementary Table 2**), which clustered into nine glial cell types and 80 neuronal populations with unique marker genes (see **Fig. 1b-e**). To keep a consistent naming convention, neurons were grouped into three major categories of glutamatergic (Glu), GABAergic (GABA) and cholinergic neurons (ChAT). Of the 80 neuronal cell populations, 48 were not part of our previous AP-centric DVC atlas (**Supplementary Figure 1a**).

**Figure 1 –.**
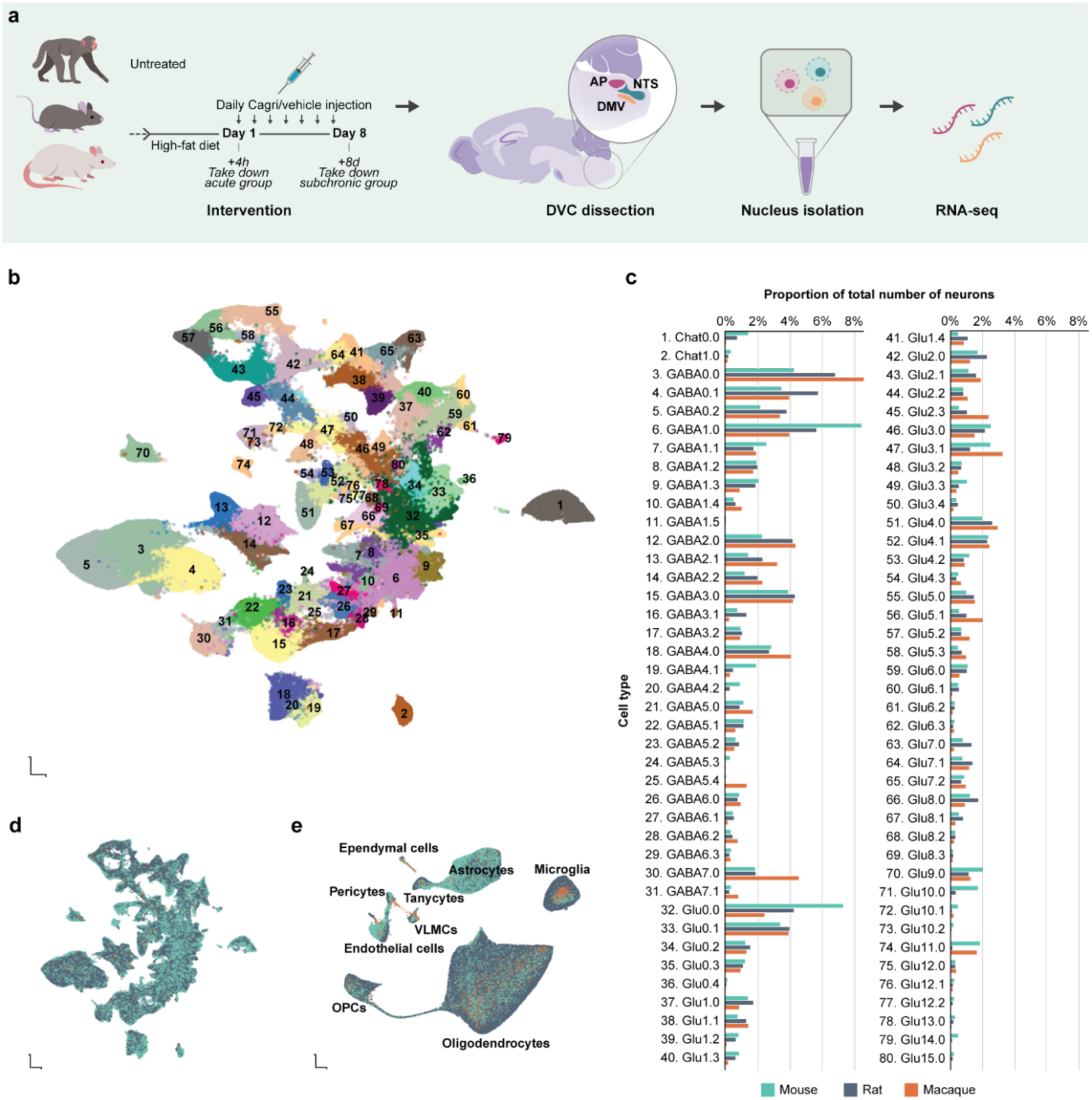
Single-cell transcriptomics cross-species atlas of the DVC. **a**, Overview of the in vivo study with acute and subchronic treatment of diet-induced obese (DIO) rats and mice, plus an addition of untreated macaques. **b**, UMAP plot of all 262,226 neurons colored by cell population. **c**, Proportions of each neuronal cell population contributing to the total number of neurons per species are plotted as corresponding bar graphs. **d**, UMAP plot of the neurons colored by species (macaque=3,404; mouse = 138,747; rat=120,075). **e**, UMAP of the 319,447 glial cells colored by species (macaque=31,056; mouse=122,382; rat=176,009). Abbreviations: UMAP, uniform manifold approximation and projection; VLMCs, vascular and leptomeningeal cells.

In contrast to our 2021 study, which utilized AP-centric dissections of the DVC and consequently excluded portions of the DVC and neighboring regions, this study employed a broader dissection strategy encompassing the entire DVC. As a result, we identified 48 new subpopulations, including several novel DVC-specific cell types (n=8) and additional cell types residing outside the DVC (n=28), with the remaining cell types not mapped to a specific region, significantly expanding the cellular diversity captured in this atlas.

Our updated atlas contained a comparable number of cells from mice and rats (251,129 vs. 296,084), and 34,460 non-human primate cells. Across all the neuronal populations from these three species, we recovered a median number of transcripts (unique molecular identifiers) of >2,500 and a median number of uniquely detected genes of >1,400, which is >1.5-fold higher compared to our previous AP-centric mouse DVC atlas and other published studies (**Supplementary Table 3**) ^11^. The proportion of major cell types varied across species: neurons comprised 55.3% of mouse cells, 40.6% of rat cells, and 9.9% of non-human primate cells. Conversely, oligodendrocytes made up 61.2% of non-human primate cells, 43.6% of rat cells, and 24.6% of mouse cells, consistent with the reported negative correlation between brain mass and relative proportions of neurons to glial cells across species ^18^.

### Spatial mapping of the rat DVC

The DVC is a spatially heterogeneous region, where cell populations expressing the same receptor(s) can respond differentially to circulating signals depending on their location. To assign cell populations to their spatial locations within the AP, NTS, and DMV, we profiled 13 DVC sections from four lean Sprague-Dawley rats using single-molecule fluorescence *in situ* hybridization provided by the Resolve Biosciences Molecular Cartography™ platform using a manually curated panel of 100 genes (**Fig. 2a, Supplementary Table 4**). Due to the longitudinal orientation of the NTS along the anterior-posterior axis, we decided to include sagittal sections close to the midline as well as standard coronal sections. All brain sections were manually region-annotated according to an anatomical atlas ^19^ and ordered by their relative position within the brain, resulting in 11 coronal sections covering a representative cross-section of the rat DVC and two sagittal sections capturing the medial rat DVC (**Fig. 2b**). Using the combined signal of the DNA-binding stain DAPI and mature RNA staining with poly(T) hybridization, the effective position and size of each cell in the section was estimated and each transcript falling within this segment was assigned to that cell (**Fig. 2c**). This resulted in the segmentation of 98,998 cells across 13 sections with a median of 196 transcripts and 33 genes per cell (**Fig. 2d, Supplementary Figure 2**), which allowed a distinction of major neuronal and glial cell types (**Supplementary Figure 3a-b**). Based on the digitized spatial dataset, we then quantified transcripts in each cell and mapped cells onto our cross-species snRNA-seq atlas (prediction score>0.6).

**Figure 2 –.**
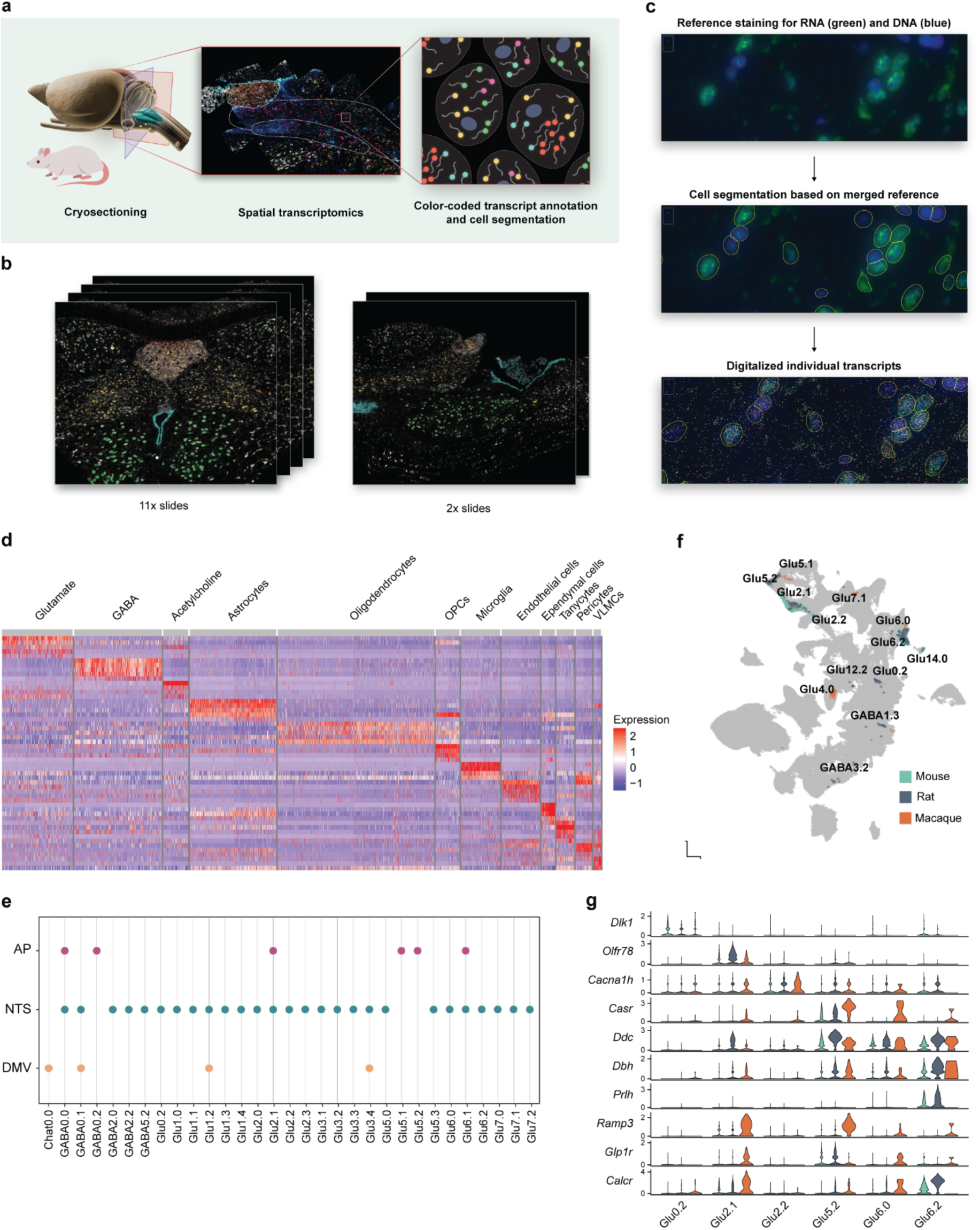
Spatial transcriptomics atlas of the rat DVC. **a**, Overview of experimental setup for sagittal and coronal sectioning of the rat DVC and schematic cell segmentation with transcript annotation. **b**, Overview of raw transcriptomic data with detected transcripts plotted over scanned brain sections. Eleven coronal sections cover a wide range of the DVC, accompanied by two sagittal sections near the midline. **c**, Detailed view of the cell segmentation process: DAPI staining (blue) and poly(T) staining (green) give a background signal for DNA and RNA in the tissue. The channels are merged, and the resulting stack undergoes automated cell segmentation. The transcripts detected through Molecular Cartography™ are then plotted over it and assigned to each segmented cell. **d**, Heatmap of gene expression across major cell types identified in the spatial rat atlas. **e**, Neuronal cell populations identified in the single-cell atlas mapped to the annotated DVC regions in the spatial rat atlas. Only cell types that map with more than random confidence (>0.6 prediction score) to the DVC are included and plotted by their statistical spatial enrichment in either the NTS, DMV, or AP. **f**, All neuronal cell population that express calcitonin receptor (*Calcr*) with high confidence (ESµ >0.8) in at least one species are plotted over the single cell atlas UMAP. **g**, Selected marker genes for each *Calcr*-expressing cell population selected based on ESµ>0.8 in each of the three species, and non-zero expression in >0.20 of the cells in the given cell population in each species. *Prlh*, was only measured in mice and rats due to few cells in the rhesus macaque samples. Abbreviations: UMAP, uniform manifold approximation and projection; VLMCs, vascular and leptomeningeal cells.

We anatomically mapped the snRNA-seq-derived neuron populations that were covered by at least 10 cells within our tissue samples (68 of 80 populations; 85%). Of these, 68 spatially defined populations, 33 (48.5%) mapped within the DVC, 28 (41.2%) resided in neighboring brainstem areas, and seven (10%) did not significantly enrich in any specific anatomical area (**Fig. 2e, Supplementary Figure 3d, Supplementary Table 5**). We have made the combined cross-species snRNA-seq and spatial atlas accessible through an interactive, publicly available user interface (See URLs).

Cagrilintide acts via CALCR (alone or in combination with RAMPs), hence understanding the cellular targets for cagrilintide requires identifying and analyzing *Calcr*-expressing cells. We used the CELLEX tool to compute combined expression specificity values (*ES*_μ_; range (0,1); low to high cell population specificity; **Supplementary Table 6**) across individual metrics for each species ^20^. Thirteen of the 70 neuron cell populations expressed *Calcr* (ES_μ_>0.8) in at least one species (**Fig. 2f**). We focused on the subset of six DVC cell populations that contained *Calcr* expression in mice or rats (**Fig. 2g**). Of these six neuron groups, a greater proportion of Glu2.1, Glu5.2, and Glu6.2 neurons expressed *Calcr* (and at higher levels) than Glu0.2, Glu2.2, and Glu6.0.

Two of the six rodent *Calcr* neuronal populations (Glu2.1 and Glu5.2) co-expressed *Ramp3* (a pattern conserved in non-human primates), permitting the formation of the amylin receptor, Amy3R – a heterodimeric complex of the CALCR and RAMP3 (**Fig. 3a**). We mapped the frequency of all six *Calcr* neuron populations to the subdivisions of the DVC and along the anterior-posterior axis (**Fig. 3b,c**; **Supplementary Figure 4a,b**). Both rodent *Calcr/Ramp3*-coexpressing populations mapped to the AP, although Glu2.1 was distributed along the AP/NTS border in rats and contained cells in the NTS as well as the AP (**Fig. 3d**).

**Figure 3 –.**
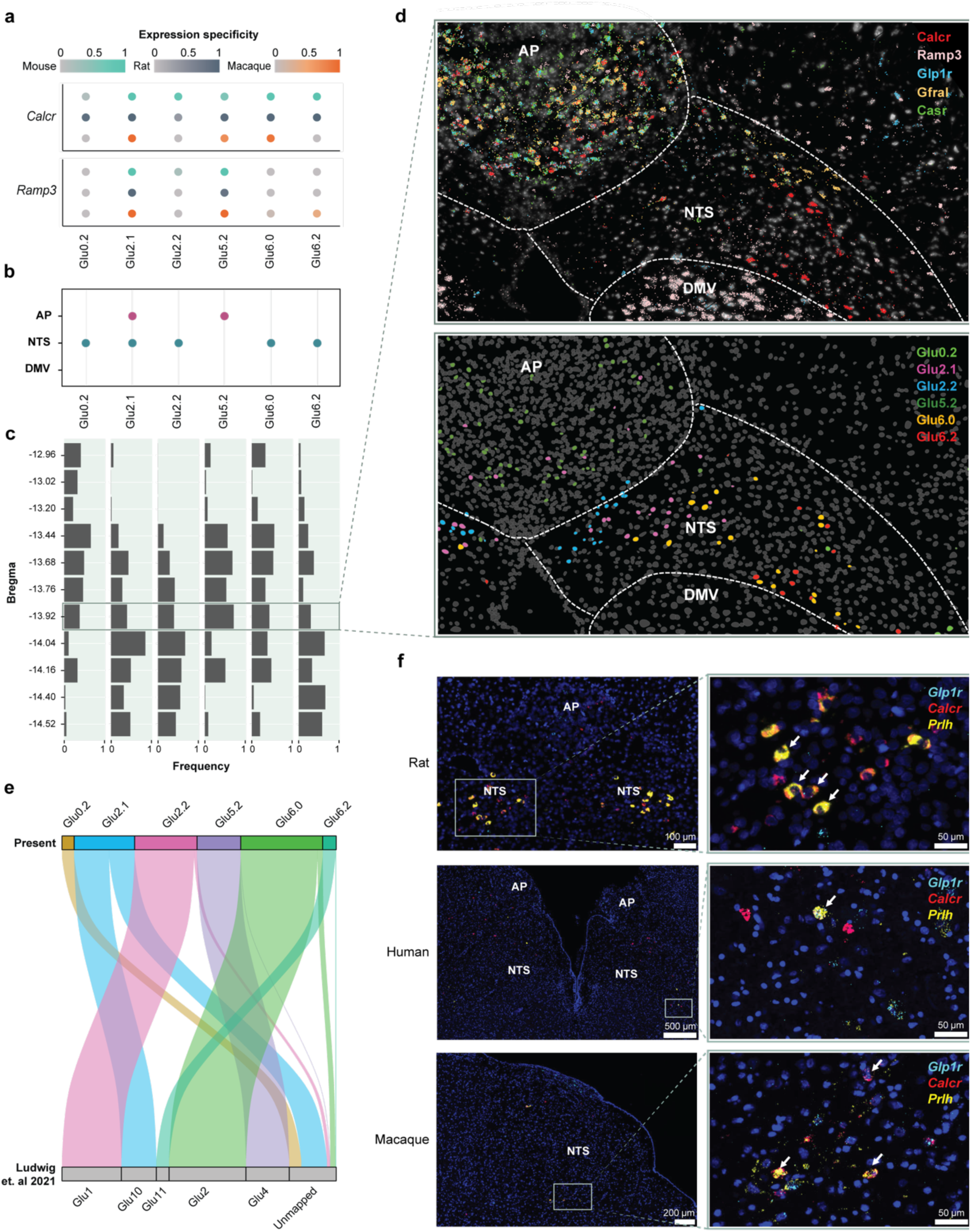
Conservation of cell populations across mice, rats, and macaques, and spatial location of *Calcr*-positive cell populations in the rat. **a**, Expression of *Calcr* and *Ramp3*, the components of the heterodimeric amylin 3 receptor, across *Calcr*-expressing neuronal cell populations. Only cell populations expressing *Calcr* with high confidence (ESµ>0.9) in the rat and/or mouse and successfully mapped to the rat spatial atlas are included. **b**, Significant presence of each cell population across the NTS, DMV and AP after Fisher’s exact test. **c**, Frequency of each given cell population through varying bregma levels based on data derived from the 11 coronal sections in the rat spatial atlas. **d**, Upper panel: DAPI+PolyT scan with detected *Calcr, Ramp3, Glp1r, Casr* and *Gfral* transcripts plotted to represent the raw spatial transcriptomics data (Resolve Polylux plugin to ImageJ). Lower panel: Segmented, digitalized and annotated spatial dataset representing the previously identified six cell population annotations transferred from the single-cell atlas. Plot from the RShiny Ratlas (Coronal section Bregma −13.92). **e**, Alluvial diagram mapping Calcr-positive cells to our previous DVC atlas. **f**, RNAScope multiplexed in situ hybridization data for Prlh, Calcr and Glp1r in rat (left panel), rhesus macaque (middle panel), and human (right panel) brain tissue shows that the defining features of the Glu6.2 cell type can be found in the ventral NTS of rodents and primates. Abbreviations: AP, area postrema; *Calcr*, calcitonin receptor; DMV, dorsal motor nucleus of the vagus; *Ramp3*, Receptor activity modifying protein 3; *Glp1r*, glucagon-like peptide-1 receptor, *Gfral*, GDNF family receptor alpha like; *Casr*, Calcium-sensing receptor; NTS, nucleus of the solitary tract; NHP, non-human primate.

The Amy3R-containing Glu2.1 AP/NTS and Glu5.2 AP populations could be distinguished by their expression of *Olfr78* or *Casr*, respectively, across species. Glu2.1 AP/NTS Amy3R/*Olfr78* neurons mapped to the previously reported AP *Calcr/Ramp3* population from mice (GLU10) ^11^. Glu5.2 AP Amy3R/*Casr* neurons mapped to the previously reported *Glp1r/Gfral/Casr*-expressing mouse AP population, which can be segmented into three subpopulations in mice (GLU4a-c from ref. ^21^; **Supplementary Figure 1b**); Glu5.2 AP Amy3R/*Casr* mapped specifically to GLU4a, which is marked by *Tnfsf11b* and expresses very little *Calcr* in mice ^11,16,21^. Glu5.1, which did not contain *Calcr* in any species, mapped to GLU4b-c.

Glu2.2 cells were located in the medial and intermediate NTS, while Glu6.0 cells were scattered in the intermediate and ventral NTS (**Fig. 3d**), as were Glu6.2 neurons (the previously identified NTS *Calcr/Prlh* neuron population in mice (GLU11)) that also contains *Glp1r*-expressing cells ^10,11,22^; **Fig. 3e**). Glu0.2, which expressed *Calcr* mainly in the rat, was confined in the ventrolateral NTS. We were able to show that Glu2.1, Glu5.2, and Glu6.0 neurons expressed *Calcr* in the rhesus macaques as well as in rodents (*ES*μ=>0.97, *ES*μ=>0.57, and *ES*μ=>0.90, respectively). Conversely, we were unable to effectively determine potential *Calcr* expression in rhesus macaque Glu6.2 neurons because only four rhesus macaque Glu6.2 neurons were detected. Thus, to confirm the location and conservation of these cells across rodents, non-human primates and humans, we used RNAScope *in situ* hybridization to investigate co-expression of the unique Glu6.2 marker *Prlh* with *Calcr* and *Glp1r* in brain sections from rats, non-human primates and humans (**Fig. 3f**). This analysis identified *Prlh/Calcr/Glp1r* co-expressing cells in the ventral NTS from these species, as predicted based on our cross-species DVC atlas and rat spatial transcriptomics data.

A few *Calcr*-expressing neuron populations (including Glu2.1 AP/NTS Amy3R/*Olfr78*, Glu5.2 AP Amy3R/Casr, and Glu6.2 NTS *Calcr/Prlh* cell groups) contained *Glp1r*-expression, suggesting that some neurons of these classes might directly respond to both GLP1RAs and amylin analogs. Our spatial transcriptomic data revealed substantial *Calcr/Glp1r*-coexpression in rat Glu5.2 AP Amy3R/*Casr* cells, but not in Glu2.1 AP/NTS Amy3R/*Olfr78* and Glu6.2 NTS *Calcr/Prlh* cells (**Supplementary Fig. 3c**), indicating that many Glu5.2 AP Amy3R/*Casr* neurons may respond to both amylin analogs and GLP1RAs, while individual Glu2.1 AP/NTS Amy3R/*Olfr78* and Glu6.2 NTS *Calcr/Prlh* cells presumably respond only to one or the other.

### Engagement of NTS Glu6.2 *Calcr/Prlh* neurons by cagrilintide in rats but not in mice

To identify potential cellular mediators of cagrilintide action, we used expression changes of the immediate early gene *Fos* and its downstream target genes (together referred to as the *Fos* regulon) as a proxy for neuronal activation to identify *Calcr*-expressing neuronal populations activated during cagrilintide treatment. We found that both acute (4 hour) and subchronic (7 day) cagrilintide treatment promoted *Fos* regulon in Glu2.1 AP/NTS Amy3R*/Olfr78* cells in the rat (*p_adj_*=0.037 and *p_adj_*=0.026, respectively; **Fig. 4a**). Interestingly, subchronic cagrilintide treatment also increased *Fos* regulon expression in the Glu6.2 NTS *Calcr/Prlh* population in rats (*p_adj_*=0.026; **Fig. 4a**). We detected no increased *Fos* regulon expression in Glu5.2 AP Amy3R*/Casr* neurons under any conditions. These results suggest acute cagrilintide treatment in rats activates Glu2.1 AP/NTS Amy3R*/Olfr78* cells while subchronic treatment activates both Glu2.1 AP/NTS Amy3R*/Olfr78* and Glu6.2 NTS *Calcr/Prlh* cells; both cell populations respond to cagrilintide in rats but not in mice.

**Figure 4 –.**
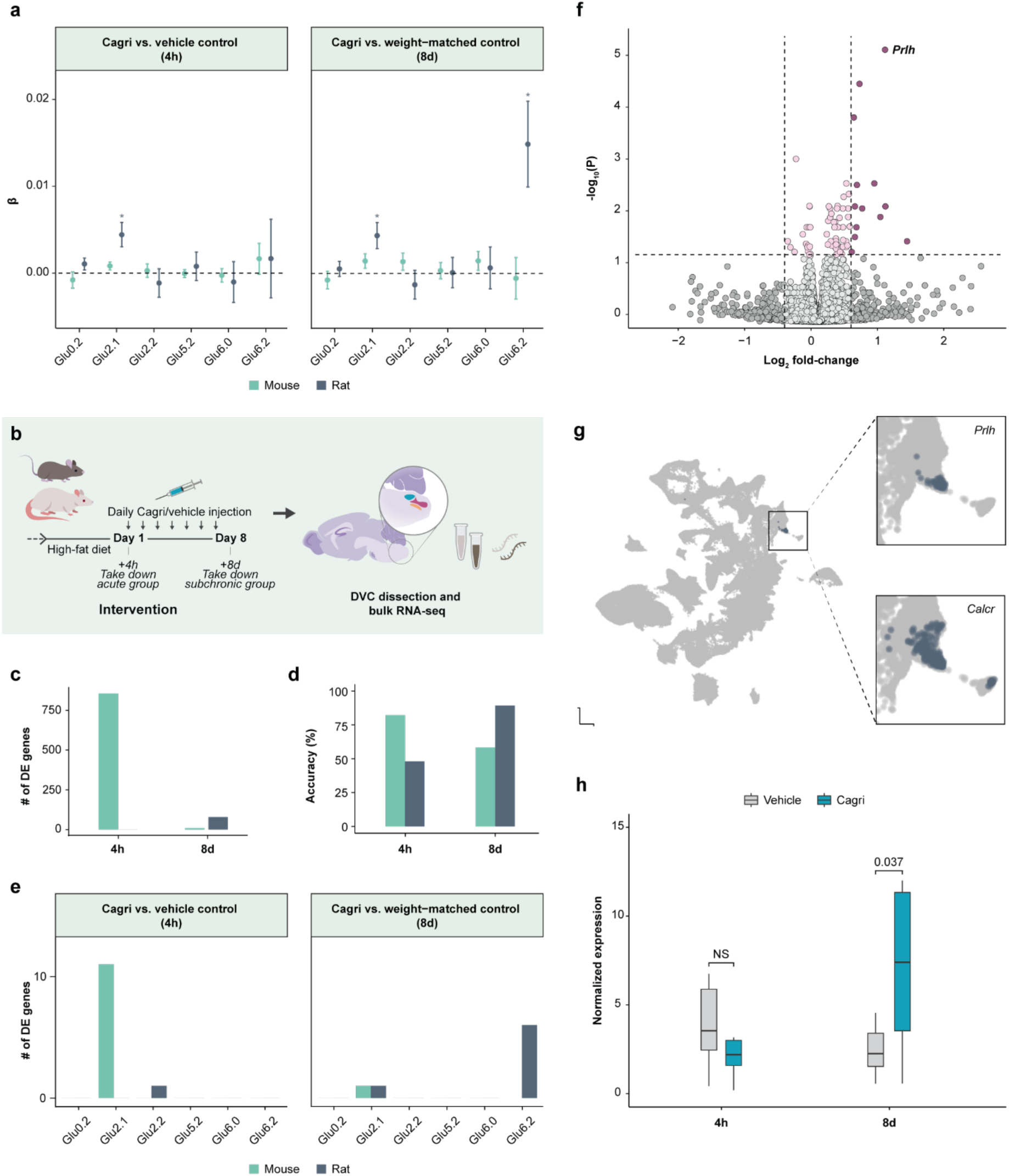
Cagrilintide activates NTS Prlh-expressing neurons and Prlh gene expression in rats but not in mice. **a**, Change in activity of Fos-regulated genes in *Calcr* neurons in mice and rats between exposure to cagrilintide and control administration after single-nucleus RNA-seq of the DVC. Only the six cell populations expressing *Calcr* in mice and rats are shown. Values are the mean estimate ± SE. *P<0.05 versus vehicle; linear mixed-effects model with sample and *Calcr* expression included as covariates (*n* = 6 mice per group, 10-11 rats per group), followed by an BH-adjusted two-tailed least-squares means t-test. **b**, Overview of the rat and mouse in vivo studies used for bulk RNA-seq. **c**, Number of differentially expressed genes between cagrilintide-administered and control animals in the acute (4h) and subchronic (8d) groups in the bulk RNA-seq experiment. **d**, Accuracy of regularized logistic regression classifiers to discriminate between cagrilintide-administered and control animals. e, Cell population specific differential gene expression in mice and rats under the same experimental paradigm of cagrilintide treatment vs. vehicle. **f**, Volcano plot for subchronically cagrilintide-administered and weight-matched control rats. Red color indicates a log_2_ fold-change >0.5 and BH-adjusted DESeq2 P < 0.05. Five data points with a log_2_ fold-change of >3 were omitted from this graph. **g**, Feature plot of normalized *Prlh* expression in the extended DVC atlas (Fig. 1b, cluster 61) marking a specific cell population. *Calcr* feature plot of the magnified UMAP region is shown as well. **h**, Expression of *Prlh* in Glu6.2 (*Calcr*/*Prlh*) neurons from rats in the acute and subchronic treatment groups (*n* = 10-11). *P < 0.05 versus vehicle; two-tailed Wald test (DESeq2). Abbreviations: DE, differential expression; DVC, dorsal vagal complex; *Prlh*, prolactin-releasing hormone; UMAP, uniform manifold approximation and projection.

To increase our ability to identify differentially expressed genes following cagrilintide treatment, we treated another set of DIO rats and mice with acute or subchronic cagrilintide (as above) and subjected the resulting DVC tissue to bulk RNA-seq (**Fig. 4b**). For each species, we performed differential expression analysis to identify transcriptional changes between cagrilintide-treated and control animals. Whereas acute treatment with cagrilintide induced 856 differentially expressed genes in mice (*p_adj_<0.05*), we detected no differentially expressed genes in acutely treated rats (although 423 genes were nominally significant). Conversely, while subchronic cagrilintide treatment detectably changed the expression of only 10 genes in mice, we identified 79 differentially expressed genes in subchronically-treated rats (1,519 genes at *p_adj_*<0.5, **Fig. 4c**).

To assess the level of cagrilintide-induced transcriptional alterations by a complementary approach, we trained regularized logistic regression classifiers to discriminate between cagrilintide-treated and control animals of each species based on gene expression. We repeatedly predicted the treatment group label of one held-out sample, which was not used for training the model, and computed how well the expression data allowed us to correctly classify a sample as originating from a cagrilintide-treated or control animal. In the acute study, the classifiers were able to discriminate between cagrilintide-treated and control mice at an accuracy of 82.1% (23/28 correctly classified mice; empirical *P_adj_*=0.016) but not in rats (accuracy=48.1%). Conversely, in the subchronic study, the classifiers could not accurately predict whether mice were treated with cagrilintide or vehicle (accuracy=58.3%), while the classifier exhibited an accuracy of 89.3% for rats (25/28 correctly classified rats; empirical *P_adj_*=0.016; **Fig. 4d**). These results suggest that subchronic cagrilintide robustly alters gene expression in the rat (but not mouse) DVC, despite only 79 genes having significantly altered expression.

We also conducted differential gene expression analysis across all *Calcr*-expressing cell populations from our snRNA-seq study (**Fig. 4e**). This analysis confirmed the robust regulation of gene expression in Glu2.1 AP/NTS Amy3R*/Olfr78* neurons by acute cagrilintide treatment in mice. Although Glu2.1 AP/NTS Amy3R*/Olfr78* cells displayed a few differentially expressed genes in both rats and mice treated with subchronic cagrilintide, rat Glu6.2 NTS *Calcr/Prlh* neurons exhibited the most differentially expressed genes, although they were not regulated in mice (**Fig. 4e; Supplementary Figure 5**). As for *Fos* regulon expression, this analysis identified no differentially expressed genes in Glu5.2 AP Amy3R*/Casr* neurons.

While bulk RNA-seq suffers from the lack of single-cell resolution, it has a high sensitivity to detect gene expression changes. Performing differential gene expressions analysis of our rat bulk RNA-seq revealed that *Prlh* was the gene most significantly differentially expressed and upregulated upon subchronic cagrilintide treatment in rats (log_2_ fold-change=1.01, *p*_adj_=5.6×10^−6^), although it was unchanged in mice (**Fig. 4f**). Glu6.2 NTS *Calcr/Prlh* neurons represent the only DVC cell population to express *Prlh*; feature plotting of normalized *Prlh* and *Calcr* expression on the single-cell atlas UMAP space shows the distinct expression of *Prlh* in one area, which at the cell population-level co-localizes with *Calcr*-expressing cells (**Fig. 4g**; compare **Figure 1b**, cluster 61). Examining *Prlh* gene expression in the snRNA-seq data for rat Glu6.2 NTS *Calcr/Prlh* neurons in isolation validated the increased expression of *Prlh* in these cells following subchronic cagrilintide treatment (log_2_ fold-change=1.60, *P*=0.038; **Fig 4h**).

Together, the results of our two different transcriptomics approaches applied to two independent *in vivo* studies suggest the activation of Glu2.1 AP*/NTS* Amy3R/*Olfr78* neurons by cagrilintide in mice and rats and demonstrate the activation and regulation of gene expression (including increased *Prlh* expression) in Glu6.2 NTS *Calcr/Prlh-*expressing neurons by subchronic cagrilintide treatment in rats but not in mice. We found no evidence for the regulation of Glu5.2 AP Amy3R*/Casr* neurons by cagrilintide under these treatment conditions. While it is possible that cagrilintide activates Glu5.2 AP Amy3R*/Casr* neurons at a time point that we have not examined, the failure of subchronic cagrilintide to alter the *Fos* regulon or other aspects of gene expression suggests that these neurons may not participate in long-term cagrilintide responses. Furthermore, either the subchronic activation of Glu2.1 AP/NTS Amy3R*/Olfr78* neurons might mediate the long-term suppression of food intake in rats, but not mice, or the increased activity and/or expression of *Prlh* in Glu6.2 NTS *Calcr/Prlh* neurons (which occurs in rats, but not mice) might mediate the long-term suppression of feeding and body weight by cagrilintide.

### AP *Calcr* neuron activation rapidly decreases food intake, but does not alter food intake or body weight over the long term

To determine the ability of Glu2.1 AP/NTS Amy3R/*Olfr78* and Glu5.2 AP Amy3R/*Casr* neurons to mediate the long-term suppression of body weight in rats, we injected an adeno-associated virus (AAV) to cre-dependently express hM3Dq-mCherry into the AP of our *Calcr^Cre^* knock-in rat model ^11^, permitting the chemogenetic activation of AP *Calcr* neurons in aggregate by treatment with the otherwise inert ligand CNO in these Calcr^AP-Dq^ rats (**Fig. 5a**). We confirmed the restriction of hM3Dq-mCherry expression to the AP by examining the distribution of mCherry-immunoreactivity, discarding any animals that had misplaced injections and/or leak of virus into neighboring regions (**Fig. 5b**). CNO treatment in the Calcr^AP-Dq^ rats promoted the accumulation of FOS immunoreactivity throughout the AP and neighboring NTS (**Fig. 5c**), demonstrating the CNO-dependent activation of the transduced AP neurons in these animals and suggesting that AP *Calcr* neurons project to and activate neurons in the NTS.

**Figure 5 –.**
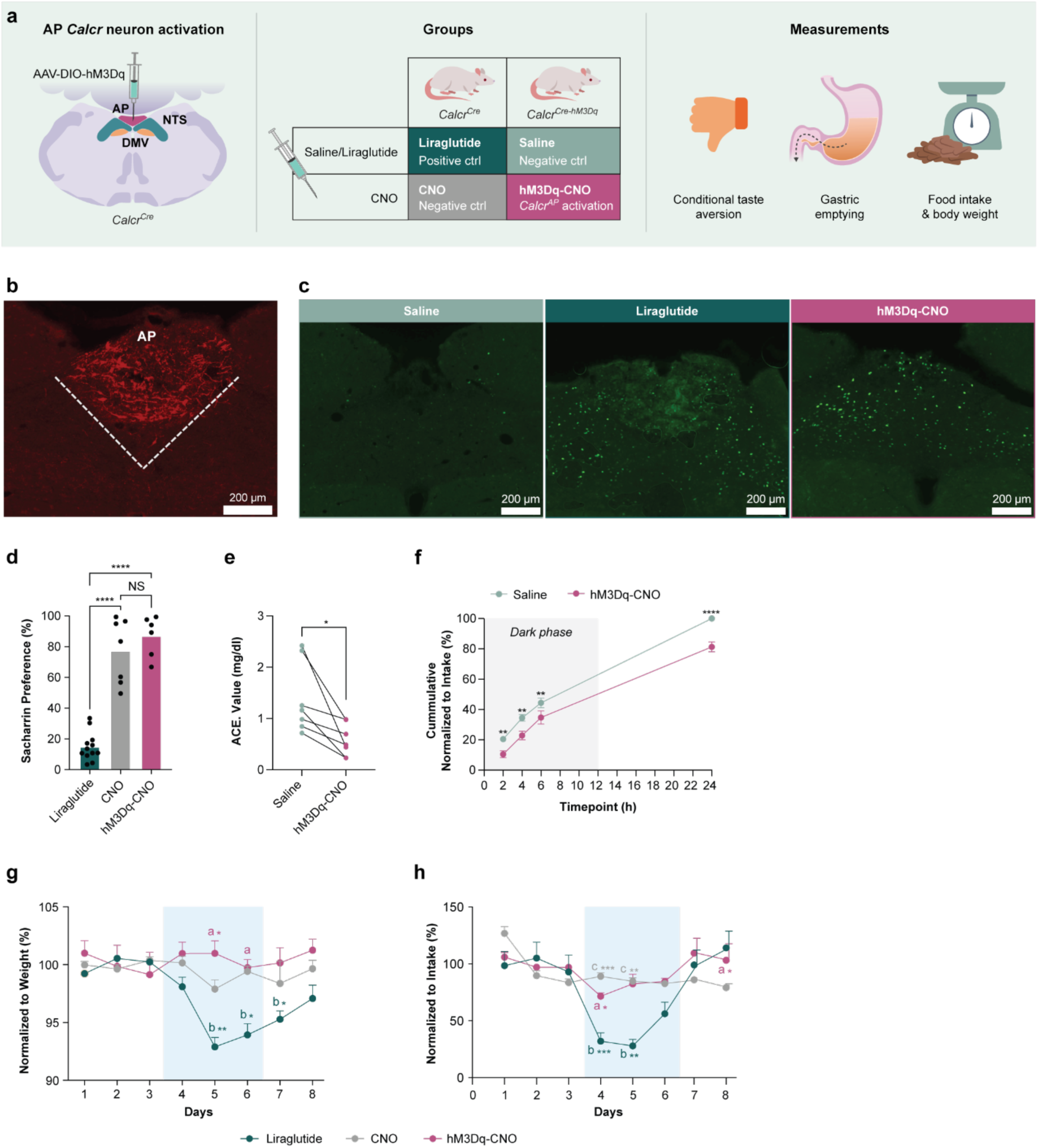
Artificial activation of rat Calcr^Cre^ neurons with hM3Dq-mCherry DREADDs by CNO administration recapitulates acute treatment effect of CALCR agonists but has no long-term effect on body weight or food intake. **a**, Schematic showing stereotaxic injections of activating DREADD virus into the AP. Behavioral paradigms included a between-subjects experimental design in non-surgery animals positive control (liraglutide) and a negative control (CNO). A within-subjects experimental design for surgery animals comparing saline vs. CNO. Measures included preference in a conditioned taste aversion study, acetaminophen levels in the blood (an indicator of gastric emptying), food intake, and body weight. **b**, Representative image of AP showing DsRed-stained neurons to confirm viral injection site (scale bar, 200 um). **c**, Visualization of cFOS immunoreactivity in the rat DVC after saline, liraglutide, and CNO treatment on AP Calcr^Cre^-DREADD rats. Representative of 4-7 animals per condition). **d**, Effect of CNO treatment in DREADD rats on conditioned taste aversion (n_liraglutide_=12, n_CNO_=7, n_hM3Dq-CNO_=6). **e**, Within-subjects experimental design: AP Calcr^Cre^-hM3Dq rats were treated with saline vs. CNO, and acetaminophen levels were measured to evaluate gastric emptying (n=7 per group). **f**, Within-subjects experimental design: The effect of CNO-induced activation of AP *Calcr*-expressing neurons was assessed by measuring 24h cumulative food intake normalized to individual intake (n=8 per group). **g, h**, Between-subjects experimental design: Normalized body weight and food intake after CNO treatment of Calcr^Cre^-DREADD rats compared to liraglutide and saline treatment of Calcr^Cre^ rats (n_liraglutide_=14, n_CNO_=8, n_hM3Dq-CNO_=8). One-way ANOVA with values in mean ± s.e.m **(d)**, Paired T-Test **(e)**. Two-way ANOVA with Šídák’s multiple comparisons test with values in mean ± s.e.m **(f)**. *p*<0.05; \*\**p*<0.01; \*\*\**p*<0.001; \*\*\*\**p*<0.001, two-tailed (d-f). Linear mixed-effects model with sample as covariate, followed by Tukey’s multiple comparisons test for CNO vs Calcr^Cre^ hM3Dq-CNO ^a^\**p*<0.05; \*\**p*<0.01; \*\*\**p*<0.001; \*\*\*\**p*<0.001, liraglutide vs Calcr^Cre^ hM3Dq-CNO ^b^\**p*<0.05; \*\**p*<0.01; \*\*\**p*<0.001; \*\*\*\**p*<0.001, liraglutide vs CNO ^C^\**p*<0.05; \*\**p*<0.01; \*\*\**p*<0.001; \*\*\*\**p*<0.001, two-tailed **(g, h)**. Abbreviations: ACE, acetaminophen; AP, area postrema; CNO, Clozapine-N-oxide; DREADD, Designer Receptors Activated Only by Designer Drugs; hM3Dq, human M3 muscarinic DREADD, Gq-coupled signaling).

We also found that a single CNO injection delayed gastric emptying and (consistent with our previous findings) decreased food intake in Calcr^AP-Dq^ rats by over 50% at the beginning of the dark cycle (**Fig. 5e-f**). This decrease in cumulative food intake continued over subsequent measurements, although the suppression of food intake relative to control waned at later times, such that the CNO-stimulated decrease in food intake was only approximately 15% lower than that of controls by the end of 24 hours (**Fig. 5f**). Hence, the suppression of food intake by the acute activation of AP *Calcr* neurons is strongest at shorter times following activation, similar to activation of those neurons with *Calcr* ligands at the beginning of food consumption in the dark phase ^7^.

Despite the strong acute suppression of food intake by the activation of AP *Calcr* neurons, we found that CNO treatment failed to promote conditioned taste avoidance (a measure of the perception of gastrointestinal malaise) in Calcr^AP-Dq^ rats (**Fig. 5d**), consistent with previous findings in mice ^16^.

To determine the ability of AP *Calcr* neurons to suppress long-term food intake and body weight, we treated Calcr^AP-Dq^ rats with CNO twice per day over three days and compared their food intake to that of control animals treated with CNO or the GLP1RA liraglutide (which strongly decreases food intake across species; **Fig. 5g-h**). While liraglutide (used here as a positive control for AP neuron activation) significantly suppressed food intake and body weight over the 3-day treatment period in control rats, CNO promoted only a small decrease in food intake in Calcr^AP-Dq^ rats during the first 24 hours of treatment (as in **Fig 5f**) and failed to decrease food intake over subsequent days of treatment or to decrease body weight at all. Thus, activating AP *Calcr* neurons in rats rapidly decreased food intake and gastric emptying, but was insufficient to mediate the long-term suppression of food intake and body weight. Hence, the long-term responses to cagrilintide in rats requires the engagement of neural systems other than the Glu2.1 AP Amy3R*/Olfr78* and Glu5.2 AP AMyR*/Casr* populations. Together with our foregoing analysis of cagrilintide-stimulated transcriptomic changes in the DVC (see Figures 3 and 4), this finding suggests a role for Glu6.2 NTS *Calcr/Prlh* neurons in the response to subchronic cagrilintide.

## Discussion

In this study, we leveraged single-nucleus and spatial profiling techniques to generate a comprehensive spatially resolved caudal hindbrain single-cell atlas and to identify DVC cell populations and molecular pathways that mediate the effects of cagrilintide. By analyzing more than 680,650 cells across the mouse, rat and rhesus macaque brainstem, we constructed an atlas covering 80 neuronal cell populations, 31 of which reside within the DVC. Among these, we identified six transcriptionally conserved cell populations that express *Calcr* in rodents, including Glu2.1 AP/NTS Amy3R*/Olfr78* neurons, Glu5.2 AP Amy3R*/Casr neurons*, Glu6.2 NTS *Calcr/Prlh* neurons, and three other lowly *Calcr*-expressing neuronal populations in the NTS. While we detected no cagrilintide-mediated transcriptional changes in Glu5.2 AP Amy3R*/Casr* neurons under the conditions tested, we found that cagrilintide activates and/or alters gene expression in the Glu2.1 AP/NTS Amy3R*/Olfr78* population in both rats and mice. Furthermore, we found that acute cagrilintide treatment activates Glu6.2 NTS *Calcr/Prlh* neurons and increases their expression of *Prlh* only in rats. This finding aligns with stronger and more persistent food intake and weight loss responses to cagrilintide in rats than mice, suggesting Glu6.2 *Calcr/Prlh* neurons in NTS to be important roles for long-term cagrilintide-mediated weight loss. However, AP *Calcr* populations in the rat were not affected. Indeed, we found that activating AP *Calcr* neurons acutely decreases gastric emptying and food intake in rats but fails to mediate the long-term suppression of food intake and body weight. Hence, the prolonged suppression of food intake and body weight by cagrilintide treatment in rats might be mediated by Glu6.2 NTS *Calcr/Prlh* neurons.

Although our earlier and other single-cell atlases of the AP and adjacent NTS have been published ^11,15,16,23^, the narrow dissection parameters used in these previous studies did not ensure complete coverage of the entire DVC. Additionally, previous studies have not examined the conservation of cell populations and receptor expression across species (including rhesus macaques), nor did they provide detailed information about the location of each cell population. The present study accomplishes each of these tasks, provides an easily accessible cross-species single-cell and spatial resource of the caudal brainstem (see URLs), and determines the DVC cells responsible for the action of amylin analogs like cagrilintide.

Because our cross-species atlas contains a larger proportion of cells from mice and rats than rhesus macaques, variation present in the rodent data might bias the cell populations identified. Furthermore, some neuron populations (e.g., Glu6.2 NTS *Calcr/Prlh* neurons) made up less than 1% of the total number of neurons. Hence, having more cells and animals per treatment group could improve the detection of differential gene expression. While the prominent transcriptional response in the bulk RNA-seq data mitigated this to some extent, these population-averaged measures may mask gene expression alterations in rare populations.

Across the three species, we identified six sets of conserved neurons that express *Calcr*. We and others had assigned the Glu2.1 Amy3R*/Olfr78* population exclusively to the AP in the mouse ^16,24^, however, in the rat this population is distributed along the AP/NTS border, with some cells on either side of the border. Additionally, we found one set of exclusively AP *Calcr*-expressing neurons (Glu5.2 Amy3R*/Casr* neurons that co-express *Glp1r*), and four *Calcr*-expressing NTS populations (Glu0.2, Glu2.2, Glu6.0, and Glu6.2 *Calcr/Prlh* neurons). Of these populations, Glu2.1 AP/NTS Amy3R/*Olfr78*, Glu5.2 AP Amy3R/*Casr,* and Glu6.2 NTS *Calcr/Prlh* neurons expressed *Calcr* in a greater number of cells and at higher levels than the other populations. While all these populations were conserved across species, the expression of *Calcr* in Glu0.2 cells appeared to be unique to the rat. Silencing *Calcr* neurons in the rat NTS was lethal, while silencing mouse NTS *Calcr* neurons promotes weight gain on HFD ^10^. Therefore, the rat-specific Glu0.2 neurons likely play roles required for survival; it is not clear what these roles might be, however.

In contrast to previous results from our group and other investigators that identified AP-residing mouse *Glp1r/Gfral/Casr*-expressing Glu5.2 neurons ^16,24^, we now show that this cell population also expresses *Calcr* across species (albeit at such low levels and in so few cells in the mouse that it may not be functionally relevant in this species). Although the activation of AP *Glp1r* neurons in aggregate promotes a conditioned taste avoidance in mice ^16^, we found that activating *Glp1r*-expressing Glu5.2 AP Amy3R*/Casr* neurons together with Glu2.1 AP/NTS Amy3R/*Olfr78* neurons produces no such effect. Hence AP *Glp1r*-expressing Glu5.2 Amy3R/*Casr* neurons appear to promote non-aversive responses. This finding thus suggests that the other AP *Glp1r* neuron population (i.e., Glu5.1 cells) mediates the aversive suppression of food intake.

Interestingly, not all *Calcr*-expressing neurons exhibited a transcriptional response to treatment with cagrilintide. Most surprisingly, we observed no increase in *Fos* regulon or overall gene expression by acute or subchronic cagrilintide in Glu5.2 AP Amy3R*/Casr* neurons. The low expression of *Calcr* in mouse cells may underlie the lack of cagrilintide-mediated effects in this species. However, the strong expression of *Calcr* in rat Glu5.2 AP Amy3R*/Casr* neurons and the widespread cagrilintide-mediated activation of the transcriptional *Fos* signature in the rat AP suggest it is unlikely that cagrilintide fails to activate these cells in rats. While cagrilintide-induced AP FOS signal peaks between one to two hours after treatment in mice, the induction of AP FOS by cagrilintide is delayed in the rat, extending at least until six hours (data not shown). Thus, it is possible that by collecting tissue at four hours following cagrilintide injection we missed the optimal time point to detect *Fos* regulon activity in the Glu5.2 AP Amy3R*/Casr* cells in the rat. Future experiments will have to address this gap by collecting more time points after acute treatment.

Also, in line with delayed cagrilintide-stimulated AP FOS accumulation in the rat relative to the mouse, we observed differential gene expression but not *Fos* regulon activity in mouse Glu2.1 AP/NTS Amy3R*/Olfr78* neurons after 4 hours of cagrilintide in mouse, suggesting *Fos* regulon activation had passed, or was undetectable, by this time. In contrast, *Fos* regulon activity, but not general changes in gene expression, was detected four hours after cagrilintide in rat Glu2.1 AP/NTS Amy3R*/Olfr78* neurons, consistent with a slower time course of cagrilintide responses in rats.

Although there was little effect of subchronic cagrilintide in DVC neurons in the mouse, the rat Glu6.2 NTS *Calcr/Prlh* population demonstrated not only ongoing activity of the *Fos* regulon, but many differentially regulated genes. Consistently, *Prlh*, specific to the Glu6.2 population, represented the most significantly upregulated gene in the rat DVC in response to subchronic cagrilintide. These data suggest potentially important roles for Glu6.2 NTS *Calcr/Prlh* neurons in cagrilintide action, which align with the known roles for these cells in the regulation of food intake and body weight. Activation of mouse NTS *Calcr/Prlh* neurons reduces food intake and body weight in mice and can override orexigenic signals from the hypothalamus ^6,10^. Furthermore, NTS *Prlh* overexpression prevents diet induced obesity, while, silencing NTS *Calcr/Prlh* neurons or the deletion of *Prlh* from the NTS deletion exacerbates DIO in mice ^10^. Coding mutations in GPR10, the *PRLH* receptor, are associated with severe obesity in humans ^25^.

Interestingly, *Prlh* is also one of the most significantly upregulated genes in the DVC of the AP in mice and the NTS of rats treated subchronically with semaglutide, which reduces body weight in both mice and rats ^11,26^. Hence, promoting *Prlh* signaling by Glu6.2 NTS *Calcr/Prlh* neurons may represent a common pathway for body weight reduction by multiple classes of gut peptide-mimetics. While Glu6.2 NTS *Calcr/Prlh* neurons do not express *Ramp* genes and thus do not contain Amy3R, it is possible that amylin analogs like cagrilintide can target these cells directly. Not all human and rhesus macaque *Prlh* neurons were *Calcr*-positive, however, and additional studies are needed to better understand the co-expression patterns of *Calcr, Glp1r* and *Prlh* in Glu6.2 neurons in higher order species.

In contrast to the role for Glu 6.2 NTS *Calcr/Prlh* neurons in the long-term suppression of food intake and body weight, we found that activating AP *Calcr* neurons (a combination of Glu2.1 AP/NTS Amy3R/*Olfr78* neurons and Glu5.2 AP Amy3R/*Casr* neurons) mediates only short-term effects of feeding and fails to mediate the long-term suppression of food intake and energy balance. Because previous studies showed that AP ablation reduces amylin-induced feeding suppression and FOS immunoreactivity in the NTS in rats ^5,8^, it is possible that AP *Calcr* neurons contribute to amylin analog action, even if they are not sufficient.

Our data provide a powerful and comprehensive new resource with which to understand DVC cell populations at baseline and in the context of amylin analog signaling. Specifically, our data suggest that subchronic activation and modulation of *Prlh* in NTS *Calcr* neurons is likely to mediate the prolonged weight loss response to cagrilintide. Our findings also raise new questions for the future. For instance, it will be useful to understand the regulation of Glu2.1 AP/NTS Amy3R/*Olfr78* and Glu5.2 AP Amy3R/*Casr* neurons, and to independently modulate them, enabling us to distinguish their functions. Similarly, it will be useful to precisely and independently modulate Glu5.2 AP Amy3R/*Casr* and non-*Calcr* Glu5.1 AP *Glp1r/Gfral/Casr* neurons.

Lastly, the pronounced difference in long-term response to pharmacological treatment between the two rodent models begs the question about the translational value for future experiments with Amylin analogues. We can acknowledge that the rat seems to more accurately model the treatment effect in primates, and that the residence of individual neuronal cell types in different compartment of the DVC, specifically between AP and NTS, can vary between species.

## Acknowledgement

Novo Nordisk Foundation Center for Basic Metabolic Research is an independent research centre, based at the University of Copenhagen, and partially funded by an unconditional donation from the Novo Nordisk Foundation (https://www.cbmr.ku.dk/; grant no. NNF18CC0034900). We acknowledge the Novo Nordisk, Novo Nordisk Foundation (grant no. NNF16OC0021496 to T.H.P.), the Lundbeck Foundation (grant no. R190-2014-3904 to T.H.P.), and the National Intstitute of Health (grant no. P51 OD011092 to the Oregon National Primate Research Center). M.G.M. and the University of Michigan team were supported by NIH R01 DK132008 and grants from Novo Nordisk and the Novo Nordisk Foundation Center for Basic Metabolic Research. We also acknowledge The Single-Cell Omics Platform (SCOP) at the Novo Nordisk Foundation Center for Basic Metabolic Research for technical expertise and support. Furthermore, this work was supported by a research grant from the Danish Diabetes Academy, which is funded by the Novo Nordisk Foundation (grant no. NNF17SA0031406). The R Shiny app is hosted by the Rodent Metabolic Phenotyping Platform (RMPP) at the Novo Nordisk Foundation Center for Basic Metabolic Research. Tune H. Pers takes full responsibility for the work as a whole.

## Contributions

M.Q.L., B.C., D.M.R., W.C., D.G., S.L., P.K., M.K.G., A.S., K.R., K.L.E., M.G.M., and T.H.P. designed the experiments. W.C., D.G., M.K.G., A.T., M.T., A.G., I.W., S.K., V.D., M. A.-M., P.V.S., J.H.S., and P.K. performed the animal experiments. M.H.T performed NTS surgeries and D.G. AP surgeries. B.C. and SCOP performed the single-nucleus experiments. M.Q.L. analyzed the data. M.Q.L., B.C., M.M. and T.H.P wrote the initial draft of the manuscript. All authors contributed to data acquisition, reviewed and edited the manuscript. T.H.P. is the guarantor of the manuscript.

## Data and code availability

All raw and processed snRNA-seq and spatial transcriptomics data will be made available through the Gene Expression Omnibus database. All code used to analyse the data is available at https://github.com/perslab/ludwig-coester-gordian-2025.

## URLs

Link to browser with the rat spatial data: https://cbmr-rmpp.shinyapps.io/spatial_dvc_app/

## Methods

### Mouse *in vivo* cagrilintide treatment studies

#### Mice

20-week-old diet-induced obese (16 weeks HFD) male C57BL/6J mice were obtained from Charles River France. Upon arrival to the animal unit, diet-induced obese mice were single-housed with *ad libitum* access to 60% HFD (#D12492 Research Diets) for seven weeks. Lean Sprague-Dawley rats, defined as animals maintained on a low-fat diet (LFD) with ad libitum access, were single-housed in parallel with DIO rats. These lean animals provide a physiological baseline for comparison with the altered metabolic state of DIO rats. The animals were housed under a 12-h light/dark cycle to standardize environmental conditions.

#### Randomization and dosing

Prior to study initiation, diet-induced obese mice were allocated into groups based on BW. For the acute study, mice were dosed with a single subcutaneous injection of cagrilintide (10 nmol kg^−1^), AM1213 (an Amy3R-selective amylin analog; 30 nmol kg^−1^), or vehicle 0-1 h after onset of light. The mice were euthanized by decapitation under isoflurane anesthesia 3-4 h after dosing. For the subchronic study, mice were dosed subcutaneously once a day with cagrilintide (10 nmol kg^−1^), AM1213 (30 nmol kg^−1^), or vehicle 3-4 h after onset of light for 7 days. One group of mice was weight-matched to mice dosed with cagrilintide by daily food restriction (**Supplementary Figure 5b**). The day after the final dose, 3-4 h after onset of light, animals were euthanized by decapitation under isoflurane anesthesia. The brains were excised followed by dissection of the DVC into RNAlater and stored on −80°C until further processing.

### Rat *in vivo* cagrilintide treatment studies

22-week-old diet-induced obese (16 weeks HFD) male Spraque-Dawley rats were obtained from Charles River France. Upon arrival to the animal unit, diet-induced obese rats were single-housed with *ad libitum* access to 60% HFD (#D12492 Research Diets) for four weeks. Lean rats were single housed with *ad libitum* access to LFD. The rats were housed in a 12-h light/dark cycle.

#### Randomization and dosing

Prior to study initiation, diet-induced obese rats were allocated into groups based on BW. For the acute study, rats were dosed subcutaneously with a single injection of cagrilintide (3 nmol kg^−1^) or vehicle 0-1 h after onset of light. For the subchronic study, rats were dosed subcutaneously once a day with cagrilintide (3 nmol kg^−1^) or vehicle 3-4 h after onset of light. The vehicle group was weight-matched to rats dosed with cagrilintide by daily food restriction (**Supplementary Figure 5a**). The day after the final dose, 3-4 h after onset of light, animals were euthanized by decapitation under pentobarbital anesthesia. The brains were excised followed by dissection of the DVC into RNAlater and stored on −80°C until further processing.

### Non-human primates

DVC tissue from adult non-human primate rhesus macaque monkeys (3 males, 2 females, 14-24 years old) was obtained from the Oregon National Primate Research Center (ONPRC, USA). All animals were purpose bred and born. All animal procedures were conducted in accordance with the guidelines of the Institutional Animal Care and Use Committee (IACUC) of the ONPRC. The ONPRC abides by the Animal Welfare Act and Regulations enforced by the United States Department of Agriculture. Rhesus macaques were deeply sedated with 20 mg/kg ketamine, followed by 25 mg/kg sodium pentobarbital, and then exsanguinated. Each brain was briefly flushed with heparinized saline, then removed from the skull and kept cold until dissection of the DVC area was performed. Tissue samples were rapidly frozen on dry ice and stored at −80°C until further processing.

### Rat surgeries and chemogenetics

#### Animals

All rats were housed in cages with Pure-o’Cel bedding and appropriate enrichment in the University of Michigan Vivarium. Rats were group-housed, except following surgery and during experimentation, during which time they were single-housed with extra enrichment. Food and water were provided ad libitum unless otherwise specified. The University of Michigan Animal Care and Use Committee (USA) approved procedures following the Association for the Assessment and Approval of Laboratory Animal Care and NIH guidelines.

Male Sprague-Dawley rats for snRNA-seq were purchased from Charles River Labs and fed high-fat diet (Research Diets, D03082706) from the time of receipt. Previously described *Calcr^Cre^*knock-in rats (Ludwig et al., 2021) were bred in our colony at the University of Michigan and were fed normal chow (Purina Lab Diet 5L0D).

#### Stereotaxic surgeries

Rats were handled for three days before surgeries. To perform AP intracranial microinjections, rats were anesthetized with isoflurane via an induction chamber at 3% and maintained under anesthesia at 2–2.5%, delivered via face mask. Oxygen flow was delivered at 0.8 l min−1. Carprofen (subcutaneous, 5 mg/kg) injections were administered before surgery to assist with pain and inflammation. Rats’ heads were flexed at a 90-degree angle on a stereotaxic device. A vertical incision was made onto the skin through the three muscle layers in a foramen past the occipital bone. A horizontal incision was made on the meninges, revealing the obex located caudal from the fourth ventricle. Stereotaxic (Model 942 Kopf with digital display console) surgeries were performed using a Hamilton syringe (5 μl syringe 800 series, 33-gauge small hub RN needle) for injections. Adult *Calcr^Cre^*rats of both sexes were used for AP injections of AAV8-hSyn-DIO-hM3Dq-mCherry (University of Michigan Viral Vector Core), coordinates were zeroed in at the center of the AP, and the virus was delivered unilaterally at a volume of 0.5 µL at coordinate Z= −0.3 mm. For NTS injected animals body weight and *ad libitum* food intake were monitored weekly for 5 weeks post-surgery.

#### DREADD-mediated activation of rat AP Calcr-neurons

*Calcr^Cre-hM3Dq^* rats were tested no sooner than 21 days post-surgery to allow for adequate viral expression. For short-term feeding and gastric emptying studies, we performed crossover studies in which order of exposure to vehicle (0.9% sodium chloride, IP.; Hospira) or CNO (Hello Bio; HB6149) was randomized. Each animal received a washout period of at least 5 days between studies. For short-term food intake, animals received IP injections 30 minutes before the onset of the dark cycle. Food was weighed at this time and again 2, 4, 6, and 24 hours later.

For gastric emptying, rats were fasted for 6 hours during the early part of the light cycle. The fasted rats received IP injections 5.5 hours post-fasting. Thirty minutes later (6 hours after the onset of the fast) the rats underwent an oral gavage of Ensure containing 0.1mg/g acetaminophen. We collected blood samples from a tail snip 15 minutes post oral gavage. The blood was placed into a vessel containing EDTA and plasma was collected and frozen. Plasma acetaminophen concentrations were determined using the Acetaminophen L3K assay kit (Sekisui Diagnostics, 506-30). At the end of the study, the animals were supplied with food and water *ad libitum*.

For long-term food intake, we conducted a between-subjects design in which separate groups of control *Calcr^Cre^* animals received CNO or liraglutide and *Calcr^Cre-hM3Dq^* rats received CNO. Each group underwent eight days of injections. During the first three days, all rats received saline at the onset of the light cycle and 30 minutes before the onset of the dark cycle. The following three days, *Calcr^Cre-hM3Dq^*and one set of control *Calcr^Cre^* rats received CNO (1 mg/kg, IP, b.i.d); the other set of control *Calcr^Cre^* rats (for liraglutide treatment) received saline at the onset of the light cycle and liraglutide (100ug/kg, IP) 30 minutes prior to the onset of the dark cycle. Food and body weight was measured at the time of the evening injection. All animals received saline injections (IP, b.i.d.) for the final two (recovery) days.

#### Perfusions, immunohistochemistry, and injection validation

Following the conclusion of *in vivo* studies, animals were perfused for the analysis of virus localization (and in some cases for the analysis of FOS). For FOS studies, animals were fasted overnight (16-18 hours) and then injected with CNO (1 mg/kg), saline, or liraglutide (100 µg/kg). Rats were overdosed with pentobarbital 200 mg/kg (IP) and perfused 1.5 hours (CNO or liraglutide) or 6 hours (cagrilintide) post-injection. Perfusions were carried out using a perfusion pump (Masterflex L/S Economy Variable-Speed Drive pump) with 1x PBS and 10% formaldehyde. The brain was extracted and placed in formaldehyde for 24 hours, then transferred to 30% sucrose for two days. Brain tissue was sectioned (blade from Accu-Edge—Low Profile Microtome Blade, 4689) using a microtome (Leica) at −40 °C. Sections were 35 µm thick.

All *Calcr^Cre-hM3Dq^*brains were subjected to immunofluorescent staining for dsRed to visualize injection placement. Animals with inadequate transduction of the AP and/or misplaced injections were excluded from analysis. For immunostaining, brain tissue was incubated in primary antibody (rabbit anti-FOS (Cell Signaling Technology, 2250), or rabbit Anti-dsRed (Takara, 632496) (all antibodies 1:1,000 dilution) at room temperature overnight followed by incubation with species-specific secondary Alexa Fluor anti-rabbit-488, anti-rabbit-568, (1:250, Invitrogen, A-11008). Sections were mounted on glass, superfrost, 75mm x 25mm microscope slides (Thomas Scientific, 1170Z86), coverslipped with 22×60-1.5 microscope cover glass (Fisher Scientific, 12-541-026) and imaged on an Olympus BX53F microscope. Black and white images were pseudo-colored using Cellsens Imaging software and ImageJ.

### Single-nucleus RNA-seq

#### Generation of single-nucleus suspensions

Tissue was thawed for approximately 2 min on ice, then transferred into a 2 ml glass tissue douncer (Sigma-Aldrich, T2690) with 1 ml Nuclei EZ Lysis Buffer (Sigma-Aldrich, NUC101)) and homogenized using pestel B. Following homogenization, the suspension was incubated for 5 min on ice, filtered through a pre-wetted 40 µm mini cell strainer (PluriSelect, 43-10040-40) into a 2 ml Protein LoBind tube (Eppendorf, 0030 108.132) and centrifuged for 10 min at 1,000*g* at 4 °C. The pellet was resuspended in 250 µl Nuclei EZ Lysis Buffer and 250 µL 83% OptiPrep™ (Sigma-Aldrich, D1556), gently layered on top of 500 µL 49% OptiPrep™, in a 2 ml Protein LoBind tube, and centrifuged for 22 min at 14,000*g* at 4 °C. The isolated nuclei located at the bottom, were resuspended in 500 µl Nuclei Buffer (PBS with 1% BSA (Sigma-Aldrich, A1595), 2 mM MgCl_2_ (Sigma-Aldrich, M1028), 40 U/ml Protector RNase inhibitor (Roche, 3335402001)) in a 1.5 ml Protein LoBind tube (Sarstedt, 72.706.605) and incubated on ice for 15 min, before centrifugation for 10 min at 1,000*g* at 4 °C. The final pellet was resuspended in 100 µl Nuclei Buffer for nuclei hashing.

#### Nuclei hashing for multiplexing

To the 100 µl single-nucleus suspension, 0.5 µg TotalSeq™-A anti-nuclear hashtag with DNA-barcoded oligonucleotide (BioLegend) was added and incubated on ice for 30 min. 1 ml Nuclei Buffer was added and the suspension was centrifuged for 10 min at 1,000*g* at 4 °C. The pellet was resuspended in 200 µl Nuclei Buffer with 0.5 µg DAPI/µl (Chemometec, 910-3012).

#### Nuclei sorting

The hashed nuclei were sorted (SH800S (SONY)) using the 70 µm Sorting Chip (Sony Biotechnologies, LE-C3207). The DAPI positive nuclei were sorted into a 2 ml Protein LoBind tube with 18.8 µl RT Buffer B (10X Genomics, 2000165) for library preparation.

#### snRNA-seq library preparation

The volume of the sorted nuclei with RT Buffer was adjusted to 61.9 µl by adding Nuclei Buffer and processed using the Chromium Next GEM Single Cell 3’ Kit v3.1 (10x Genomics (1000268)), according to the manufacturer’s protocol. To generate a multiplexing Hashtag library, the BioLegend protocol (TotalSeq™-A Antibodies and Cell Hashing with 10x Single Cell 3’ Reagent Kit v3.1 (single Index) Protocol) was followed from step II-III.

### Single-nucleus RNA-seq raw data processing

#### Gene counting

Raw sequencing reads were processed using Cell Ranger v7.1.0. BCL files were converted to FASTQ files using the *mkfastq* function. If hamming distance between any adapter sequences were less than three, barcode mismatches were not allowed during the conversion. Reads were aligned to the *Mus musculus* (mm10), *Rattus norvegicus* (Rnor_6.0), or *Macaca mulatta* (Mmul_10) genome with default parameters. Gene counting was performed using both exonic and intronic reads. To remove ambient RNA, the raw gene-barcode matrix was used as input for CellBender v.0.2.2 ^28^ that was run in *cuda* mode with the parameters *expected-cells* set to 7,000, *total-droplets-included* set to 20,000, and *fpr* set to 0.1.

#### Cell counting

While Cell Ranger and CellBender were used for gene counting, an optimized framework was implemented for cell counting. Initially, the reference genomes for the three species were indexed using the salmon *index* tool v1.5.0 ^29^. Similarly, the reference for antibody alignment was created with the salmon *index* tool using sample names and HTO sequences as input. Alignment of RNA was performed with salmon alevin v1.5.0 ^30^ that was run in *sketch* mode and *library type* was defined as ISR 10x chromium v3. Alignment of HTO was likewise performed with salmon alevin run in *sketch* mode and *library type* defined as ISR protocol by setting the read geometry to read 2, position 1-15, barcode geometry as read 1, position 1-16, and UMI geometry as read 1, position 17-26.

The subsequent quantification steps were carried out within the alevin-fry framework. First, a permit list of valid cell barcodes was generated with the *generate-permit-list* function. Cell calling strategy was initially set as unfiltered mapping and quantification based on the February 2018 10x Genomics whitelist. Additionally, the expected orientation of alignments was set to both directions to capture both sense and anti-sense reads. The valid barcodes from the previous steps were then corrected using the *collate* function according to the information given in the permit list step. Finally, the reads were quantified and outputted as a count matrix using the *quant* function. The resolution strategy by which molecules were counted was set to cr-like to support splice aware alignment.

Subsequently, cell calling and doublet removal were carried out in R. Cell calling was done on the count matrix of the unspliced counts. First, barcode ranks were found with the *barcodeRanks* function from the DropletUtils v1.10.3 package ^31^. A Hartigan’s dip test was performed on the counts. If the resulting *P* value ≥ 0.05 and thus the null hypothesis of unimodality was accepted, cells above the calculated inflection were used for downstream processing. If the resulting *P* value < 0.05, a bimodal model was fitted to the counts from cells above the original inflection with the *Mclust* function from the mclust v5.4.7 package ^32^. Cell barcode ranks were then calculated again on the subset of cells assigned to cluster two (the cluster with the highest average gene counts) and cells above the re-calculated inflection were used for downstream processing. Cells overlapping between the RNA and HTO library were imported into Seurat v4.0.0 ^33^. The HTO assay was CLR-normalized and scaled. Cell barcodes were then demultiplexed into sample identity with the *HTODemux* function. The positive quantile threshold was determined by running the function with all 0.01 increments going from 0.50 to 0.99, optimizing for the maximum number of classified singlets. To remove intra-sample cell doublets, RNA counts were log-normalized, PCA was performed, and doublets were inferred from the inter-sample doublets with the *recoverDoublets* function from the scDblFinder v1.4.0 package. The filtered barcodes from the above pipeline and the gene counts from the CellBender pipeline were combined into a count matrix and used as input for downstream analysis.

### Single-nucleus RNA-seq data analysis

#### Initial processing

Downstream analysis was carried out in Seurat. Initially, snRNA-seq data for each species were processed individually. Two batches, containing cells from 16 mice, were removed due to failed library preparation, which was indicated by the following sequencing. Cells that were outliers based on UMI count or mitochondrial RNA content were identified. For the mouse snRNA-seq data, cells with a UMI count > 5 x 10^−4^, UMI count < 500, or mitochondrial RNA content > 2% were discarded. For the rat snRNA-seq data, cells with a UMI count > 3 x 10^−4^, UMI count < 500, or mitochondrial RNA content >2 % were discarded. For the macaque snRNA-seq data, cells with a UMI count > 5 x 10^−4^, UMI count < 500, or mitochondrial RNA content >0.5 % were discarded.

For the mouse and rat snRNA-seq data, batch effects were not noticeable, and cells were merged across batches using the *merge* function. Raw counts were normalized with the *SCTransform* function. For the macaque snRNA-seq data, batch effects were evident, and cells from each batch were normalized using *SCTransform* and integrated using the *FindIntegrationAnchors* and *IntegrateData* functions with *reduction* set to canonical correlation analysis (CCA) with the top 3,000 most variable genes as input. All three datasets were subjected to principal component analysis (PCA) dimensionality reduction using the *RunPCA* function with the top 3,000 most variable genes as input. To split neurons and glial cells, an initial round of clustering was performed using the *FindNeighbors* and *FindClusters* functions on the top 30 PCs with *resolution* set to 0.1. For each species, neurons and glial cells were identified using known marker genes, and above steps were repeated.

#### Integration across species

To generate a cross-species neuronal and glial cell atlas, neurons and glial cells from all three species were integrated. To increase coverage of the AP and neighboring cells, cells from an AP-centric snRNA-seq atlas published by us were included in this integration (see below for reprocessing of this dataset) ^11^.

Rat and macaque genes were mapped to mouse orthologous genes using files of orthologous genes downloaded from www.ensembl.org/biomart/martview. Glial cells and neurons were integrated using the *FindIntegrationAnchors* and *IntegrateData* functions with *reduction* set to CCA with the top 3,000 most variable genes as input. The two atlases were subjected to PCA using *RunPCA* and clustering using *FindNeighbors* and *FindClusters* on the top 30 PCs with *resolution* set to 1.

To identify low-quality clusters, for the neuronal atlas, for each dataset and cluster, the mean mitochondrial RNA content and expression of *Clic6* (marker of choroid plexus contamination) and *Mbp* (marker of oligodendrocytes; the most abundant glial cell type) were computed. For the glial atlas, for each dataset and cluster, the mean mitochondrial RNA content and expression of *Clic6* and *Slc32a1* and *Slc17a6* (markers of GABAergic and glutamatergic neurons; the most abundant neuronal cell classes) were computed. Clusters with a mean above the 95% quantile in at least three out of four datasets were marked as low-quality and discarded. In addition, for the glial atlas, to discard potentially missed doublets, clusters containing high expression of markers of different glial cell types were identified. Specifically, for each dataset and cluster, the mean expression of *Ikzf1* (microglia marker), *Slc38a5* (endothelial cell marker), *Ntsr2* (astrocyte marker), *Myt1* (OPC marker), and *Mbp* were computed. Clusters with a mean larger than one fourth of the maximum cluster mean in three out of four datasets for two different glial cell type markers were marked as doublets and removed (with the exception of *Myt1* and *Mbp* which are expressed in two developmentally similar cell types).

#### Clustering of neurons

To cluster the neurons into neuronal populations, neurons were initially split into GABAergic, glutamatergic, and cholinergic neurons based on the expression of *Slc32a1*, *Slc17a6*, and *Chat*, respectively. For each of the three broad neuronal classes, neurons were clustered using *FindNeighbors* and *FindClusters* at a *resolution* of 0.1, yielding 26 major clusters. Subsequently, each of the 26 major clusters were subclustered using the *FindSubCluster* function at a *resolution* of 0.1, yielding 83 clusters.

To identify marker genes of the 83 clusters, CELLEX v.1.2.2 was run on all three species together. For each cluster, the top marker genes were defined as the 50 genes with the highest CELLEX score while being expressed in >20% of the cells in the focal cluster.

Neighbor clusters were defined as clusters assigned to the same major cluster. Subsequently, two neighbor clusters with <10 marker genes separating them were merged. This strategy yielded a final set of 80 clusters of neuronal populations.

#### Differential expression analysis

Differential expression analysis was performed using DESeq2 v1.30.1 ^34^. For each cell population, a pseudobulk gene expression matrix was constructed by summing the raw transcript counts for cells originating from the same animal. Animal and cell population combinations with less than five cells were excluded. Cell populations with less than two pseudobulk replicates per treatment group were not included in the differential expression analysis. Neither were genes expressed in less than 10% of the cells of the focal cell population included. DESeq2 differential expression analysis was performed using the Wald test. *P* values were corrected for multiple testing using the BH method (adjusting for the number of genes).

#### SCENIC analysis

SCENIC ^35^ was run separately on the mouse and rat neurons using the python implementation v0.11.0 with default parameters. To identify whether the mouse or rat *Fos* regulon was differentially activated between cagrilintide-treated and control animals in *Calcr* neuronal populations, a linear mixed-effects model was constructed for each *Calcr* neuronal population using the regulon activity as the dependent variable, the treatment group and *Calcr* expression as independent variables, and animal as a random effect. Group contrasts were tested usings a least-squares means, two-tailed *t*-test. *P* values were corrected for multiple testing using the BH method (adjusting for the number of *Calcr* neuronal populations).

#### Reprocessing of the published AP-centric DVC atlas

To limit batch effects due to differences in raw data processing, the AP-centric transcriptomics atlas published by us ^11^ was reprocessed. Raw sequencing reads were processed using Cell Ranger v.7.1.0 as described above. The raw gene-barcode matrix was subsequently used as input for CellBender which was run with the same parameters as above. The filtered barcodes from Cell Ranger and the updated gene counts from CellBender were combined into a count matrix and used as input for downstream analysis.

Downstream analysis was carried out in Seurat. Cells with a UMI count > 1.5 x 10^−4^, UMI count < 250, or mitochondrial RNA content > 2% were discarded. Since batch effects were not noticeable, cells were merged across batches and normalized. Subsequently, PCA was performed and followed by an initial round of clustering using the top 30 PCs. The cells were then split into neurons and glial cells, and PCA were recomputed for the neurons and glial cells.

### Bulk RNA-seq

#### Library preparation

mRNA libraries were produced using the Illumina NovaSeq platform where paired-end reads (2 x 150 bp) were sequenced to a depth of approximately 2 x 10^7^ reads per sample.

#### Alignment and quantification

STAR v2.7.a3 was used for alignment and feature counting. The GRCm38 GENCODE vM23 genome and the genome were used for alignment of the mouse and rat data, respectively.

#### Outlier removal

Mouse and rat samples were subjected to vsd-transformation using DESeq2 v1.30.1 followed by PCA. Outliers were identified as being more than 3 standard deviations away from the mean PC1 or PC2.

### Bulk RNA-seq data analysis

#### Differential expression analysis

Differential expression analysis was performed with DESeq2 using the Wald test with the fraction of ribosomal RNA included as a covariate. To enable comparison between species, only orthologous genes identified in both species were included in the analysis. *P* values were corrected for multiple testing using the BH method (adjusting for the number of genes).

#### Logistic regression classification

Logistic regression classification with Lasso regularization was performed using the glmnet v4.1.1. R package. To increase speed, PCA was performed on the vsd-transformed bulk RNA-seq data and used as input for the regression analysis with the treatment group as the response variable and the PCs as the predictor variables.

To assess the performance of the regression classifier, the following steps were performed. The input data was split up into a training dataset comprising all samples except one and a test dataset comprising the held-out sample. A regression classifier was constructed on the training dataset. Leave-one-out cross-validation was applied to the training dataset for choosing the optimal regularization parameter λ through minimization of the cross-validation classification error using the *cv.glmnet* function. The trained regression classifier was subsequently applied on the test dataset to compute a test classification error. These steps were repeated so that each sample was used to compute a test classification error of a regression classifier trained on the remaining samples. A mean classification error across all samples was subsequently calculated.

#### *In vivo* data statistical analysis

Differences in body weight and food intake were assessed using a linear mixed-effects model with body weight or food intake as the response variable, treatment and day as predictor variables with interaction effects, and samples as random effects. Group contrasts were tested usings a least-squares means, two-tailed *t*-test. *P* values were corrected for multiple testing using the BH method.

### Spatial transcriptomics

#### Tissue preparation and staining

Fresh frozen rat brains from chow-fed Sprague-Dawley rats (Janvier, France) were sectioned at 12µm thickness to represent coronal and sagittal cuts of the dorsal vagal complex. The slides were then processed with a proprietary multiplexed in situ hybridisation protocol containing 100 genes (Resolve Biosciences, Germany) to yield a dataset of localized transcripts across the scanned brain areas. We went on to stain the sections for RNA to better visualize neurons for cell segmentation. Five washes with PBS were followed by 2h of biotinylated poly(T) 1:500 (Z5261, Promega) in 0.4% Triton-100 at 41°C. Another five washes with PBS were followed by 2h of Streptavidin-647 1:500 (S21374, Invitrogen) in 0.4% Triton-100 at RT. Sections were then washed again 5 times in PBS before imaging.

#### Segmentation of spatial transcriptomics

To predict segmentation of the cell nuclei, the spatial transcriptomics data was initially segmented using Cellpose v.1 ^36^. As an input, we stacked DAPI and poly(T) stainings using ImageJ ^37^. To account for the large variation of neuron sizes specifically in the hypoglossal nucleus, we used two different parameters for Cellpose segmentation and merged the different segmentations using a decision tree. The cell segmentation and the spatial transcriptomics data (transcript location and identity) were then used as input to a custom pipeline that generates a spatial Seurat object (See our GitHub repository for implementation). The Seurat object contains a regular cell-by-gene matrix as well as the spatial information.

#### Cell population label transfer from single-cell to the spatial transcriptomics data

The spatial transcriptomics data underwent quality control using the transcript count and spatial cell size as input. Afterwards the spatial transcriptomics data was normalized using the *SCTransform* function. The snRNA-data was subsetted to all common genes between snRNA data and spatial transcriptomics data. Cell population labels were then transferred from the single-cell to the spatial transcriptomics data using the *FindIntegrationAnchors* and *IntegrateData* functions with *reduction* set to canonical correlation analysis (CCA).

#### Cell population enrichment across DVC areas

To identify neuronal populations that were enriched in a specific DVC area (AP, NTS, or DMV), a contingency matrix was constructed for each combination of neuronal population and DVC area. From the contingency matrix, the odds ratio was calculated, and statistical significance was evaluated using a Fisher’s exact test with an alpha of 0.05. Correction for multiple testing was performed using Bonferroni correction (adjusting for the number of cell populations-DVC area combinations).

#### Multiplex in situ hybridization

Fluorescent in situ hybridization (FISH) was performed on formalin fixed paraffin embedded brain blocks covering the dorsal vagal complex from rat, rhesus macaques and human material. The tissue blocks were sectioned at 5 μm and mounted onto Fisher SuperFrost Plus glass slides (Fisher Scientific). Multiplex FISH was performed using the RNAscope LS multiplex fluorescent reagent kit (Advanced Cell Diagnostics, Bio-Techne, Cat #322800) together with iFluor546/iFlour594 (1:500, AAT Bioquest/VWR, Cat #AATB45035/AATB45025) and Opal690 Fluorophore Reagent pack detection (1:500, Cat #FP1497001KT, Akoya Biosciences) on a Leica BOND RX Fully Automated Research Stainer (Leica) according to the manufacturer’s instructions. The human and rhesus macaque tissue sections were pre-treated for 30 min with HIER at 95°C in ER2 (Leica) followed by 15 min protease treatment and then hybridized with human-specific probes to detect mRNA transcripts for CALCR (#483048), PRLH (#526568-C2) and GLP1R (#519828-C3) (Advanced Cell Diagnostics, Bio-Techne). The rat tissue sections were pre-treated for 15 min with HIER at 95°C in ER2 (Leica) followed by 15 min protease treatment and then hybridized with rat-specific probes to detect mRNA transcripts for CALCR (#477798), PRLH (#1301288-C2) and GLP1R (#315228-C3) (Advanced Cell Diagnostics, Bio-Techne). As negative control, ACD 3-plex Multiplex Negative Control Mix (Cat #320878, ACD/Biotechne) was used. Slides were counterstained with DAPI and then cover slipped with Prolong TM Gold Antifade mountant (ThermoFisher Scientific). Fluorescent slide scans were acquired with an Olympus VS200 slide scanner (Olympus) using a 20x (0.8 NA) air objective and 385/470;546/570;594/620;635/690 filter sets. Images were prepared with the Olympus OlyVIA software, and signal intensity levels were adjusted to match across staining/slides.

#### Tissue sources for multiplex in situ hybridization

##### Human

Tissue blocks covering caudal medulla were obtained from the Edinburgh Brain Bank (n=5, incl. males and females) in collaboration with Professor Colin Smith, following UK and DK legal and ethical guidelines. Informed consent was obtained from all participants.

##### Non-human primate

The caudal medulla from one purpose bred (for research and safety testing) male cynomolgus monkey (Macaca fascicularis) was collected at necropsy reusing vehicle animals from another study. Sourcing of these monkeys following EU Directive 2010/63.

##### Rat

Young adult lean Sprague Dawley rats (n=2: 1 male and 1 female) were anaesthetized by isoflurane and cardiac perfusion fixation was performed using a peristaltic pump, starting with NaCl for 4 min and followed by neutral buffered formalin (NBF, VWR International A/S) for 8 min. After O/N post fixation in NBF, each brain was split into 2mm slices using a brain tissue matrix and then processed to formalin-fixed paraffin embedded (FFPE) blocks.

