## Supplementary Figures 1-5 for "A Cross-Species Atlas of the Dorsal Vagal Complex Reveals Neural Mediators of Cagrilintide’s Effects on Energy Balance"

**Supplementary Figure 1**

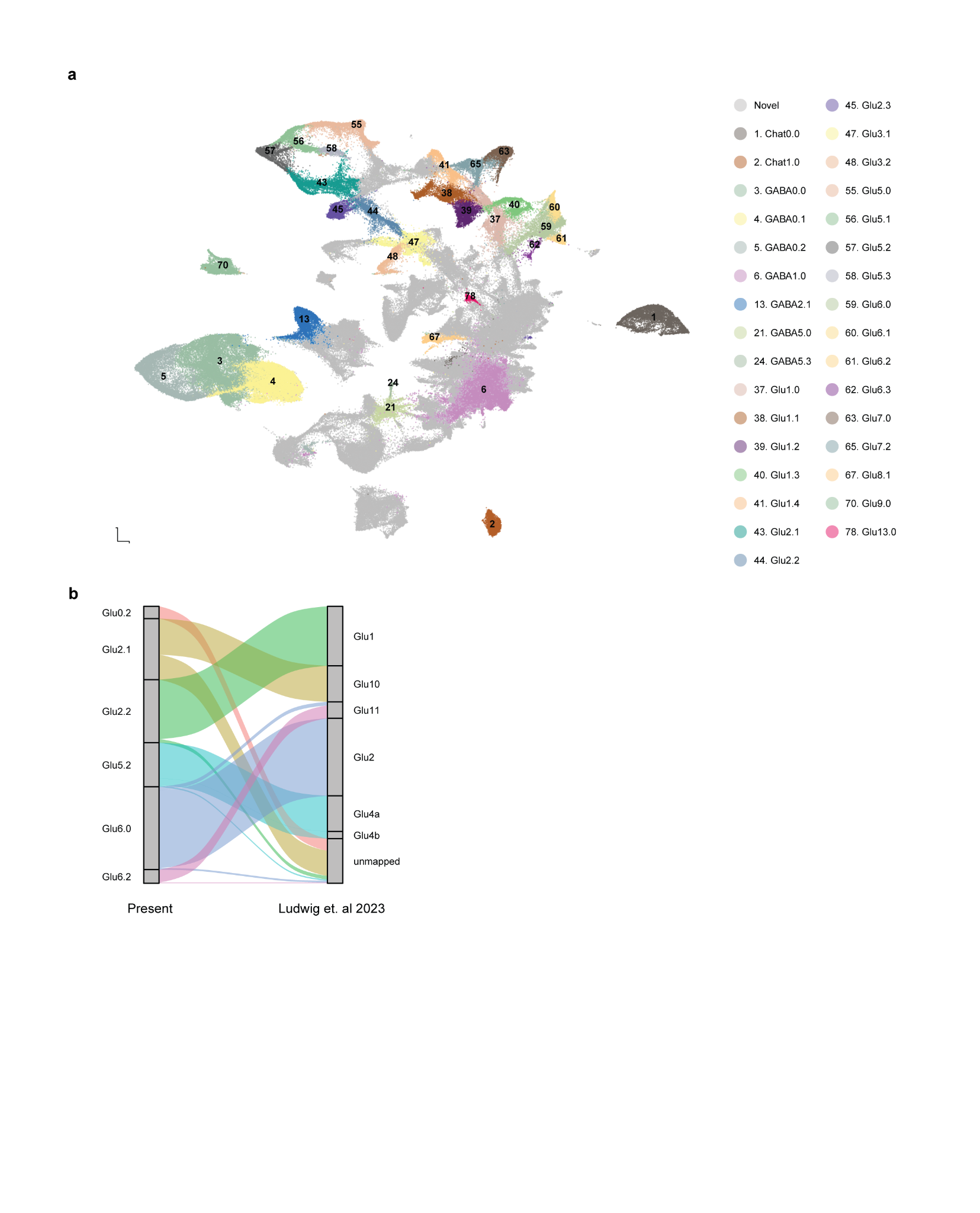

**Single-cell transcriptomics cross-species atlas of the DVC. a,** UMAP of the mouse single-cell data from (Ludwig et al. 2021) included in this atlas plotted over the entire neuronal atlas. Newly discovered regions are colored in grey, clusters from the old atlas are colored in and denoted by their new identity. **b,** Sankey plot of *Calcr*-expressing neuronal cell types in the DVC mapping to the single cell atlas from (Ludwig et al. 2023) with extended annotations (Glu4a, b). Abbreviations: UMAP, uniform manifold approximation and projection; Calcr, Calcitonin receptor.

**Supplementary Figure 2**

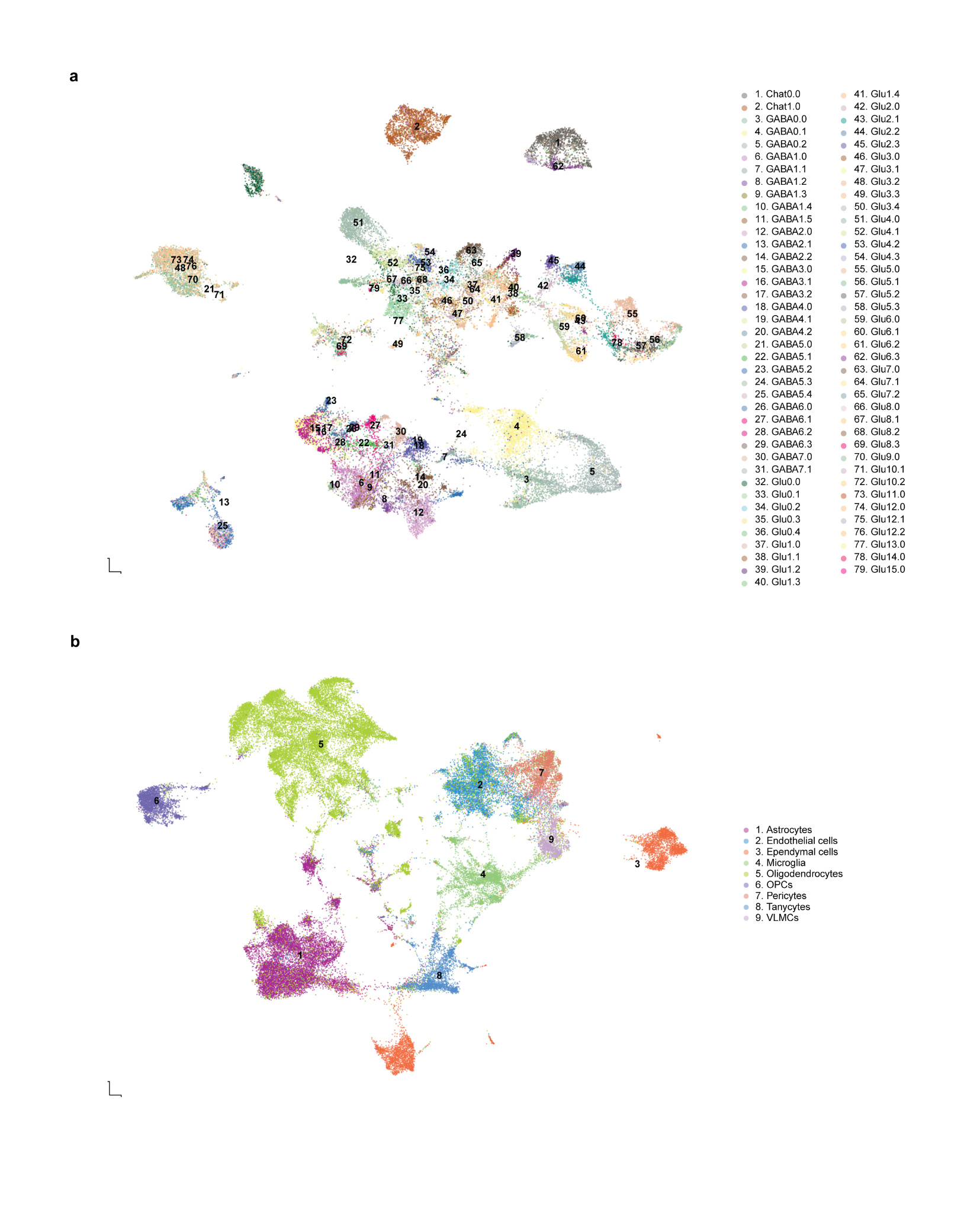

**Identified cell types in the spatial transcriptomics atlas of the rat DVC. a,** UMAP of neuronal cells identified in the digitalized spatial dataset, characterized by their assigned transcripts. **b**, UMAP of glial cell populations. All transcripts were assigned to segmented cells based on the combined signal for DNA and RNA density. All transcript counts were normalized by cell area and cells were annotated based on reference single cell data. Abbreviations: UMAP, uniform manifold approximation and projection.

**Supplementary Figure 3**

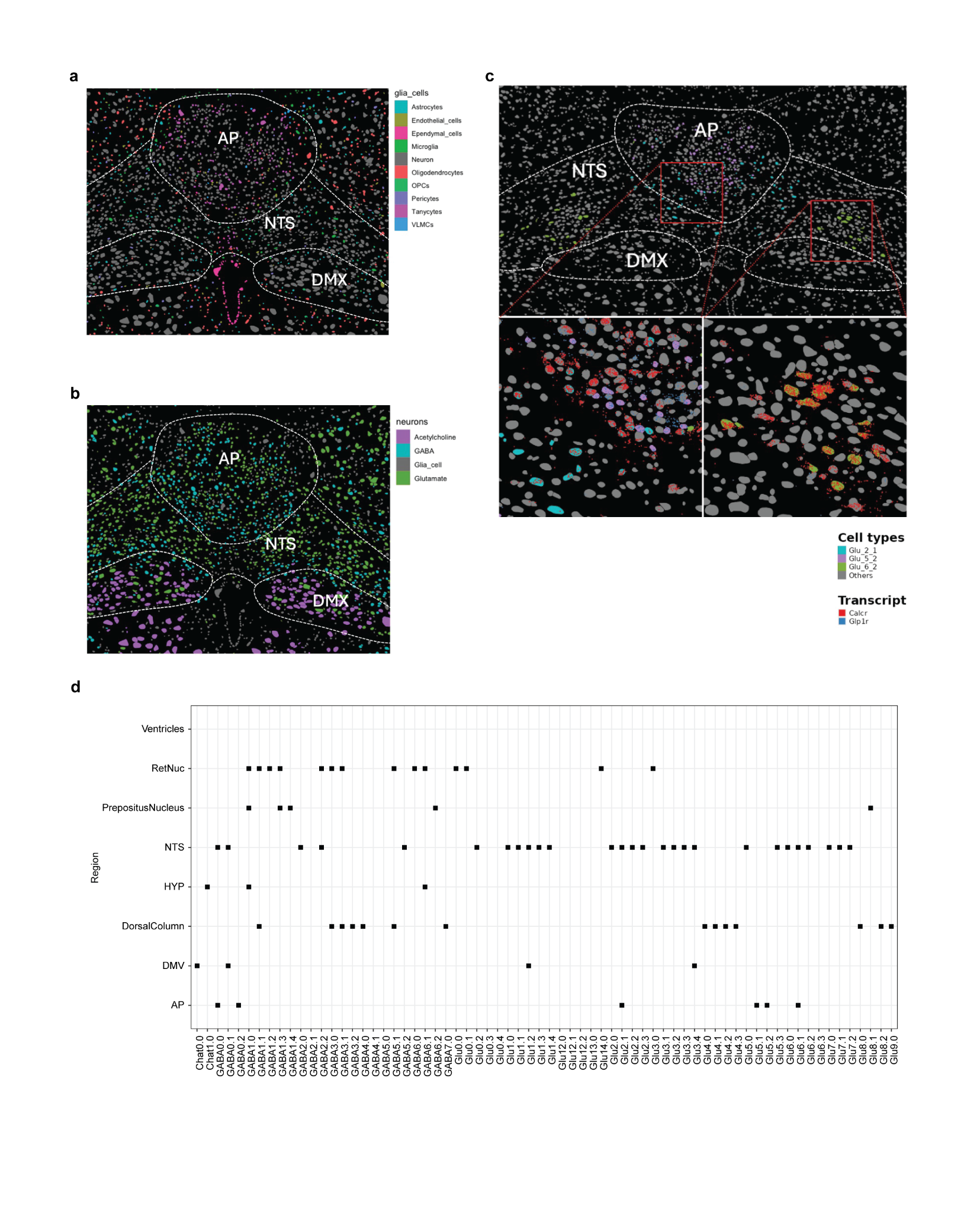

**Spatial transcriptomics atlas of the rat DVC. a,** Digitalized spatial brain section at Bregma -14.16 in the DVC with glial cells colored, neurons in grey. **b,** Neurons plotted in the same brain section, colored by major neurotransmitter classes (Acetylcholine, Chat; Gamma-aminobutyric acid, GABA; Glutamate, Glu). **c,** Digitalized spatial brain section of the DVC with three neuronal cell types highlighted (Glu2.1, Glu5.2, Glu6.2), *Calcr* and *Glp1r* transcripts plotted over the magnified regions. **d,** Expanded panel of cell type enrichment in all spatial sections by region annotation. Squares represent a significant enrichment of each cell type in the annotated hindbrain region.

**Supplementary Figure 4**

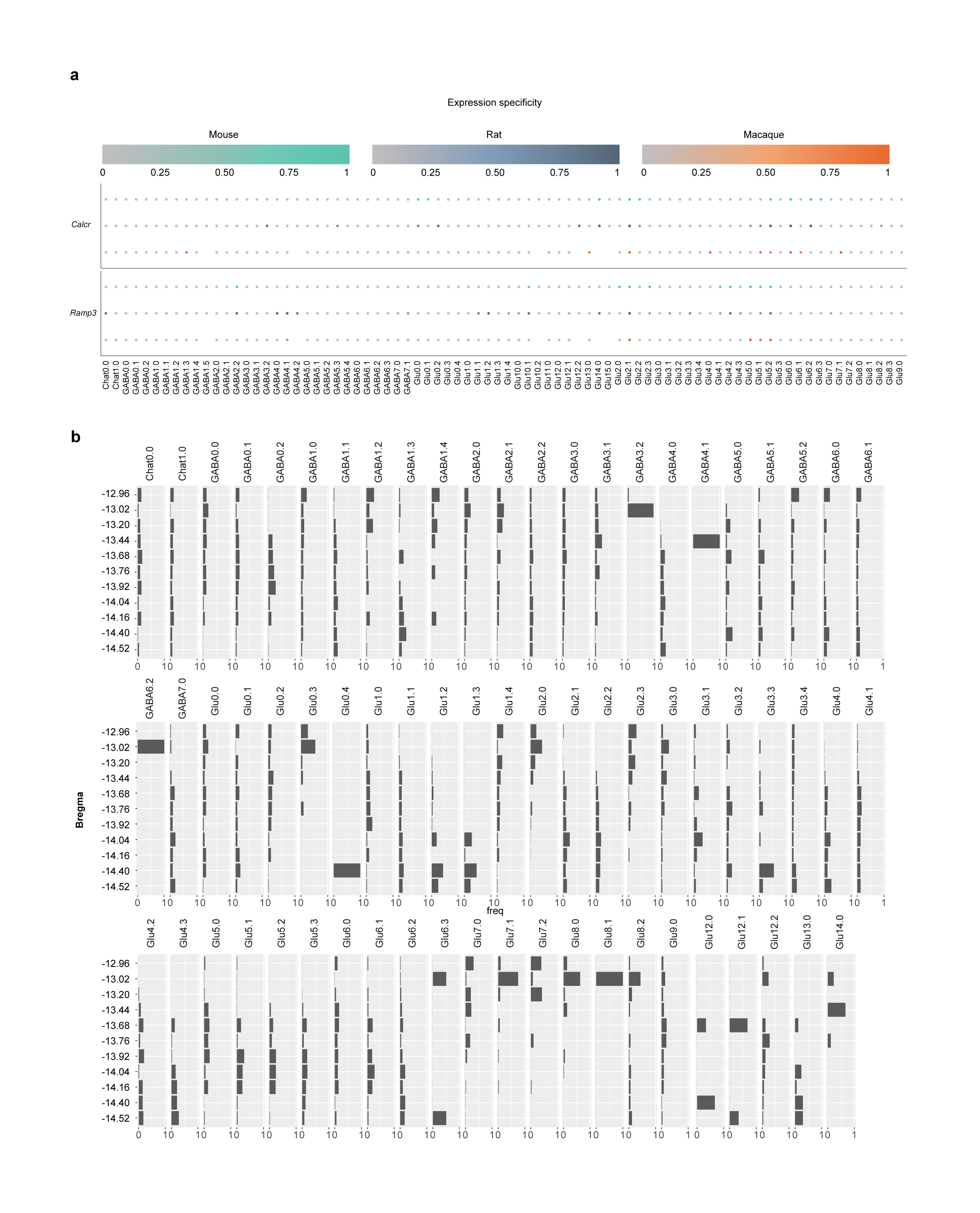

**Conservation of cell populations across mice, rats, and macaques, and spatial location of Calcr+ cell populations in the rat. a**, *Calcr* and *Ramp3* expression across neuron cell populations for each species, represented by predicted expression specificity ESu. **b**, Frequency of each neuron cell population by varying bregma levels based on data derived from the 11 coronal sections in the rat spatial atlas. Abbreviations: Calcr, calcitonin receptor; Ramp3, Receptor activity modifying protein 3.

**Supplementary Figure 5**

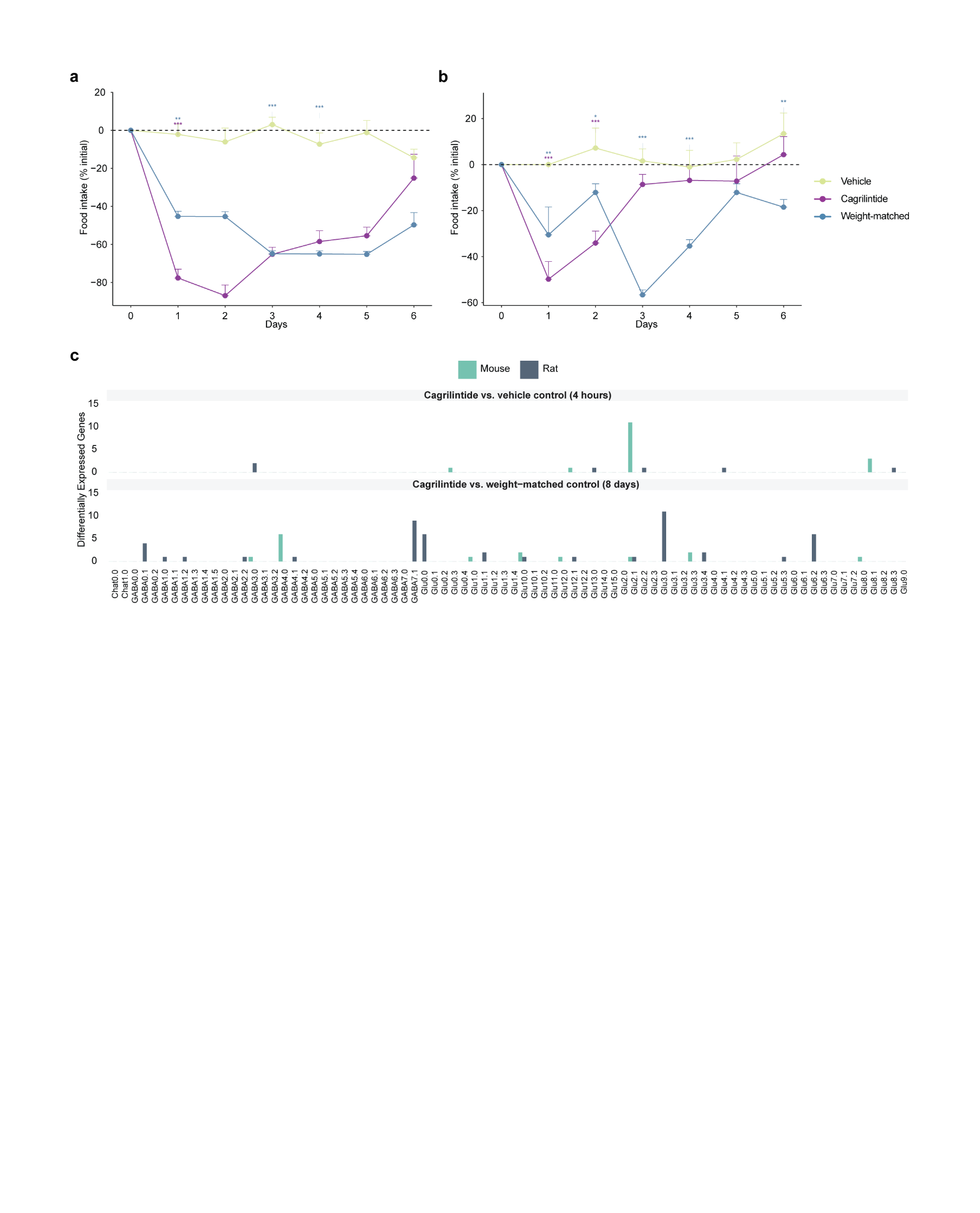

**Cagrilintide activates NTS prolactin releasing hormone-expressing neurons and Prlh gene expression in rats but not in mice. a-b,** Daily food intake relative to the initial value in rats (**a**) and mice (**b**) following cagrilintide or vehicle administration (**a** = 10 rats; **b** = 7-8 mice). Values are the mean ± s.e.m. **P* < 0.05, ***P* < 0.01, ****P* < 0.001 versus vehicle. **c,** Differentially expressed genes from bulk RNA sequencing across all cell types for acute (4h, top panel) and subchronic (8d, bottom panel) treatment for mice and rats.
